# Spontaneous neural oscillations influence behavior and sensory representations by suppressing neuronal excitability

**DOI:** 10.1101/2021.03.01.433450

**Authors:** Luca Iemi, Laura Gwilliams, Jason Samaha, Ryszard Auksztulewicz, Yael M Cycowicz, Jean-Remi King, Vadim V Nikulin, Thomas Thesen, Werner Doyle, Orrin Devinsky, Charles E Schroeder, Lucia Melloni, Saskia Haegens

## Abstract

The ability to process and respond to external input is critical for adaptive behavior. Why, then, do neural and behavioral responses vary across repeated presentations of the same sensory input? Spontaneous fluctuations of neuronal excitability are currently hypothesized to underlie the trial-by-trial variability in sensory processing. To test this, we capitalized on invasive electrophysiology in neurosurgical patients performing an auditory discrimination task with visual cues: specifically, we examined the interaction between prestimulus alpha oscillations, excitability, task performance, and decoded neural stimulus representations. We found that strong prestimulus oscillations in the alpha+ band (i.e., alpha and neighboring frequencies), rather than the aperiodic signal, correlated with a low excitability state, indexed by reduced broadband high-frequency activity. This state was related to slower reaction times and reduced neural stimulus encoding strength. We propose that the alpha+ rhythm modulates excitability, thereby resulting in variability in behavior and sensory representations despite identical input.

## Introduction

When asked to make repeated perceptual decisions, we often respond differently with each repetition, even when given the same sensory information (Rahnev & Denison, 2018; Wyart & Koechlin, 2016). We addressed the physiological sources of this variability by testing how spontaneous fluctuations in neuronal excitability, reflected by alpha oscillations (7–14 Hz), affect behavior and neural sensory representations.

While historically regarded as “noise”, spontaneous neural oscillations strongly predict neural dynamics and behavior in health (Busch et al., 2009; Grabot & Kayser, 2020; Lange et al., 2013; Romei et al., 2008b; Samaha, Iemi, et al., 2020) and disease (age-related cognitive decline: Tran et al., 2020; Voytek et al., 2015; schizophrenia: Uhlhaas & Singer, 2010; autism: Simon & Wallace, 2016). Here, “spontaneous” oscillations are operationalized as neural activity occurring without (“ongoing”) or preceding (“prestimulus”) sensory stimulation, since activity fluctuations during these moments are presumed to be endogenously generated. Strong ongoing alpha oscillations are related to a state of low excitability, as indexed by a reduction of neuronal firing (Bollimunta et al., 2008, 2011; Chapeton et al., 2019; Dougherty et al., 2017; Haegens et al., 2011; Lundqvist et al., 2020; van Kerkoerle et al., 2014; Watson et al., 2018), local field potentials (Potes et al., 2014; Spaak et al., 2012), and hemodynamic activity (Becker et al., 2011; Goldman et al., 2002; Mayhew et al., 2013). Further, states of strong prestimulus alpha oscillations predict behavioral changes, including longer reaction times (RTs; Zhang et al., 2008; Bollimunta et al., 2008; Kelly & O’Connell, 2013; Bompas et al., 2015), lower probability of reporting near-threshold sensory stimuli (Ergenoglu et al., 2004; van Dijk et al., 2008; Busch et al., 2009; Mathewson et al., 2009; Chaumon & Busch, 2014; Iemi & Busch, 2018; Limbach & Corballis, 2016; Iemi et al., 2017; Craddock et al., 2017) or phosphenes (Romei et al., 2008a; Samaha, Gosseries, et al., 2017), and reduced subjective perception (i.e., lower confidence: Samaha, Iemi, et al., 2017; Samaha, LaRocque, et al., 2020; lower visibility ratings: Benwell et al., 2017).

Despite this extensive work on alpha oscillations, the mechanisms supporting their apparent interrelation to excitability and behavior remain unclear. One proposal suggests that alpha oscillations reflect a mechanism of functional inhibition, regulating the excitability state of the neural system and thereby information processing necessary for behavior (Griffiths et al., 2019; Jensen & Mazaheri, 2010; Klimesch et al., 2007; Mathewson et al., 2011; Samaha, Iemi, et al., 2020). At the physiological level, a state of functional inhibition may be achieved, for example, via the activation of GABAergic inhibitory interneurons and/or via reduced excitatory drive (e.g., downregulation of norepinephrine or acetylcholine). According to this account, the behavioral changes associated with alpha oscillations are caused by alpha power influencing neuronal excitability. Alternatively, alpha power might affect excitability and behavior via independent mechanisms (e.g., Gundlach et al., 2020). As most studies to date report evidence for a link between alpha power and *either* excitability *or* behavior, it is currently unknown whether the relationship between alpha power and behavior is mediated by a direct excitability modulation or via an independent mechanism.

It also remains unknown whether and how the excitability modulation associated with prestimulus alpha oscillations shapes the neural representations of sensory stimuli, on which behavior depends. Using decoding as a proxy for neural representation, we compared two hypotheses of how alpha oscillations may affect how sensory information is represented in the brain. First, low excitability during strong alpha oscillations may relate to a state of reduced attentional resources (Diepen & Mazaheri, 2017; Van Diepen et al., 2019), associated with decreased neural responses (Mehrpour et al., 2020; Treue & Maunsell, 1996) but increased variability/noise (Cohen & Maunsell, 2009; Mitchell et al., 2009). The resulting lower signal-to-noise ratio may thus worsen the encoding of sensory stimuli (i.e., lower decoder accuracy). Second, low excitability during strong alpha oscillations may be related to a decrease in the neural response magnitude and variability/noise (Goris et al., 2014; Tomko & Crapper, 1974). The resulting decrease in both signal and noise may thus reduce the overall strength of neural representations (i.e., lower decoder confidence; see Methods), leaving their encoding accuracy unaffected (Samaha, et al., 2020).

Here, we tested the functional inhibition account of alpha oscillations and their effect on behavior and neural sensory representations by analyzing intracranial electroencephalography (iEEG) recordings in nine patients with medication-resistant epilepsy (N = 1044 electrodes) while they performed an auditory discrimination task with visual cues (Figure 1A; Auksztulewicz et al., 2018). iEEG enabled us to estimate ongoing neural oscillations and broadband high-frequency activity (BHA, 70–150 Hz), which is thought to reflect neuronal ensemble activation patterns, including multiunit activity (Manning et al., 2009; Nir et al., 2007; Ray et al., 2008; Rich & Wallis, 2017; Ray & Maunsell, 2011; Miller et al., 2014), dendritic processes integral to excitation of neuronal ensembles (Leszczyński et al., 2020; Suzuki & Larkum, 2017), as well as additional neuronal processes such as synaptic currents (Lachaux et al., 2012). Critical for the present study, BHA provides a reliable measure of local neuronal excitability. First, we confirmed a hallmark of the functional inhibition account: i.e., a simultaneous negative relationship between ongoing alpha power and neuronal excitability (as indexed by BHA: Potes et al., 2014; Spaak et al., 2012). We tested whether this pattern generalizes across brain areas, other frequency bands (including beta, 15–30 Hz), and processing windows (i.e., prestimulus and poststimulus), and whether it reflects a genuine oscillatory modulation, or a change in the aperiodic 1/f activity. Second, we tested whether *pre*stimulus alpha power modulates the neural response to sensory stimuli (as indexed by *post*stimulus BHA), and whether this modulation generalizes across sensory modalities (i.e., visual and auditory). Third, we tested whether the effect of prestimulus alpha power on poststimulus excitability has consequences for task performance and for decoded neural stimulus representations. Specifically, we analyzed how prestimulus alpha power influences subsequent RTs, and we used mediation analysis to directly link prestimulus alpha power with RT changes via poststimulus excitability modulation. In addition, we used multivariate pattern analysis to test whether prestimulus alpha power affects decoder accuracy and/or confidence.

**Figure 1.**
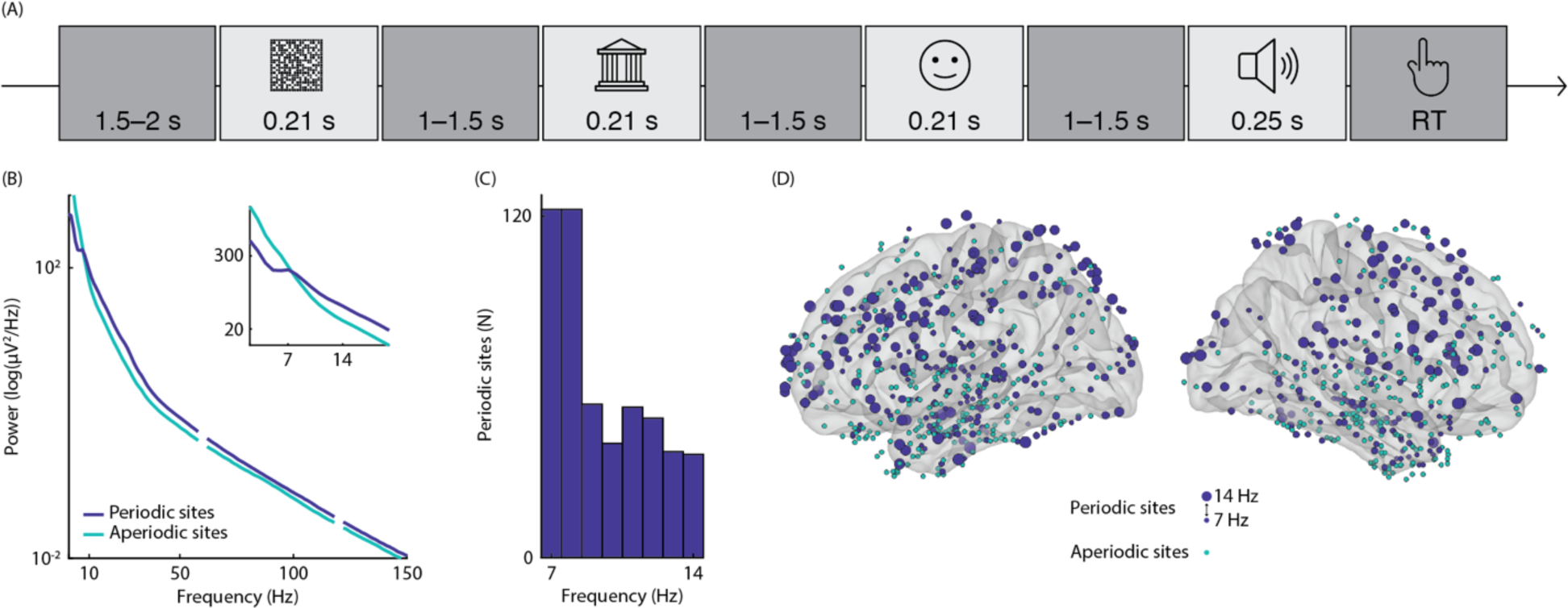
Experimental paradigm and alpha-band periodic and aperiodic activity. **A**. Schematic overview of the experimental paradigm. In each trial, participants were presented with a sequence of sensory stimuli in a fixed order: a noise image, a scene image, a face image, and a target syllable. At the end of the trial, participants reported the syllable identity (/ga/ or /pa/) via button press. **B.** Averaged power spectrum of the 1-s window before the noise image, shown separately for sites with alpha oscillations (periodic sites, in blue) and sites without a detectable alpha-band peak (aperiodic sites, in green). The inset shows the power spectrum for the frequency window of interest. **C.** Histogram of the alpha-band peak frequencies of the prestimulus power spectrum across periodic sites. **D**. Schematic illustration of the iEEG electrode coverage. Direct recordings of brain activity were obtained using intracranial electrodes implanted in 1044 sites across 9 epilepsy patients. Blue and green dots illustrate periodic and aperiodic sites, respectively; the size of blue dots indicates the alpha-band peak frequencies of periodic sites.

To preview our results, strong prestimulus oscillations in the alpha+ band (i.e., alpha and neighboring frequencies), rather than the aperiodic signal, are correlated with lower BHA, reflecting reduced excitability. This effect is observed across the brain and sensory modalities as well as before and after sensory input. Further, strong prestimulus alpha oscillations are correlated with slower RTs, and with a decrease in decoder confidence, but not accuracy. Critically, low excitability during the poststimulus window mediates the relationship between prestimulus alpha power and RTs, demonstrating a link between alpha oscillations, behavior, and excitability consistent with functional inhibition. We propose that, by modulating neuronal excitability, the ongoing alpha+ rhythm affects behavior and the strength of neural stimulus representations.

## Results

### Alpha power negatively correlates with simultaneous BHA

Based on the functional inhibition account of alpha oscillations, we hypothesized that a state of low neuronal excitability (here indexed by reduced BHA) would occur specifically during strong prestimulus alpha oscillations, rather than the aperiodic 1/f signal in the same frequency range. To test this, we sorted 1-s prestimulus epochs (i.e., window before noise image onset) into five bins based on single-trial estimates of alpha power (i.e., 7–14 Hz average) and computed the average BHA estimated in the same prestimulus window for each bin. We sorted the recording sites into “periodic” and “aperiodic” based on the presence or absence of a (local) peak within the alpha frequency range of the prestimulus power spectrum, respectively (following the methods in Haegens et al., 2014). To improve the spectral estimates, the peak detection was based on the trial-averaged data; therefore, we interpreted the peak in periodic sites as reflecting clear oscillatory (periodic) activity across trials, whereas the absence of a peak in aperiodic sites as indicating a more prominent 1/f aperiodic signal across trials (though, peaks may be present in individual trials, see Discussion). We found 525 periodic sites with alpha-band oscillatory activity (mean peak at 9 Hz, SEM=0.10), and 519 aperiodic sites with non-oscillatory 1/f activity in the alpha frequency range (Figure 1B–D; Figure 1 supplement shows periodic and aperiodic activity relative to the beta band). We hypothesized a negative correlation between prestimulus alpha power and BHA specifically for periodic sites, but not in aperiodic sites.

To test this, we used a repeated-measures mixed-effect ANOVA with BHA as dependent variable, periodic/aperiodic sites as between-unit factor (with units referring to recording sites), and alpha bins as within-unit factor. We found a significant main effect of alpha bins (F(2.18, 2274.72)=64.72, p<0.001; all ANOVAs Huynh-Feldt corrected) indicating that, across sites, BHA decreased with alpha power (bin 5 vs. bin 1: t=−13.99, p<0.001; all post-hoc comparisons Holm-Bonferroni corrected). Further, we found a significant main effect of periodic/aperiodic sites (F(1, 1042)=71.50, p<0.001), with higher BHA in periodic sites (t=8.46, p<0.001). Critically, we found a significant interaction effect between periodic/aperiodic sites and alpha bins (F(2.18, 2274.72)=53.73, p<0.001), indicating that prestimulus BHA decreased with prestimulus alpha power in periodic sites (bin 5 vs. bin 1: t=−19.51, p<0.001; Figure 2A), but not in aperiodic sites (bin 5 vs. bin 1: t=−0.33, p=1; Figure 2C). An anatomical map of this effect (Figure 2B) in periodic sites revealed a widespread BHA reduction during states of strong prestimulus alpha power across the brain. Furthermore, we replicated this interaction effect on BHA estimates in which the power at each frequency was normalized across trials (i.e., frequency-normalized BHA: F(1.94, 2021,57)=93.34, p<0.001; bin 5 vs. bin 1 in periodic sites: t=−19.59, p<0.001; in aperiodic sites: t=6.20, p<0.001), confirming that prestimulus alpha power suppressed BHA after controlling for the 1/f contribution to BHA estimates.

**Figure 2.**
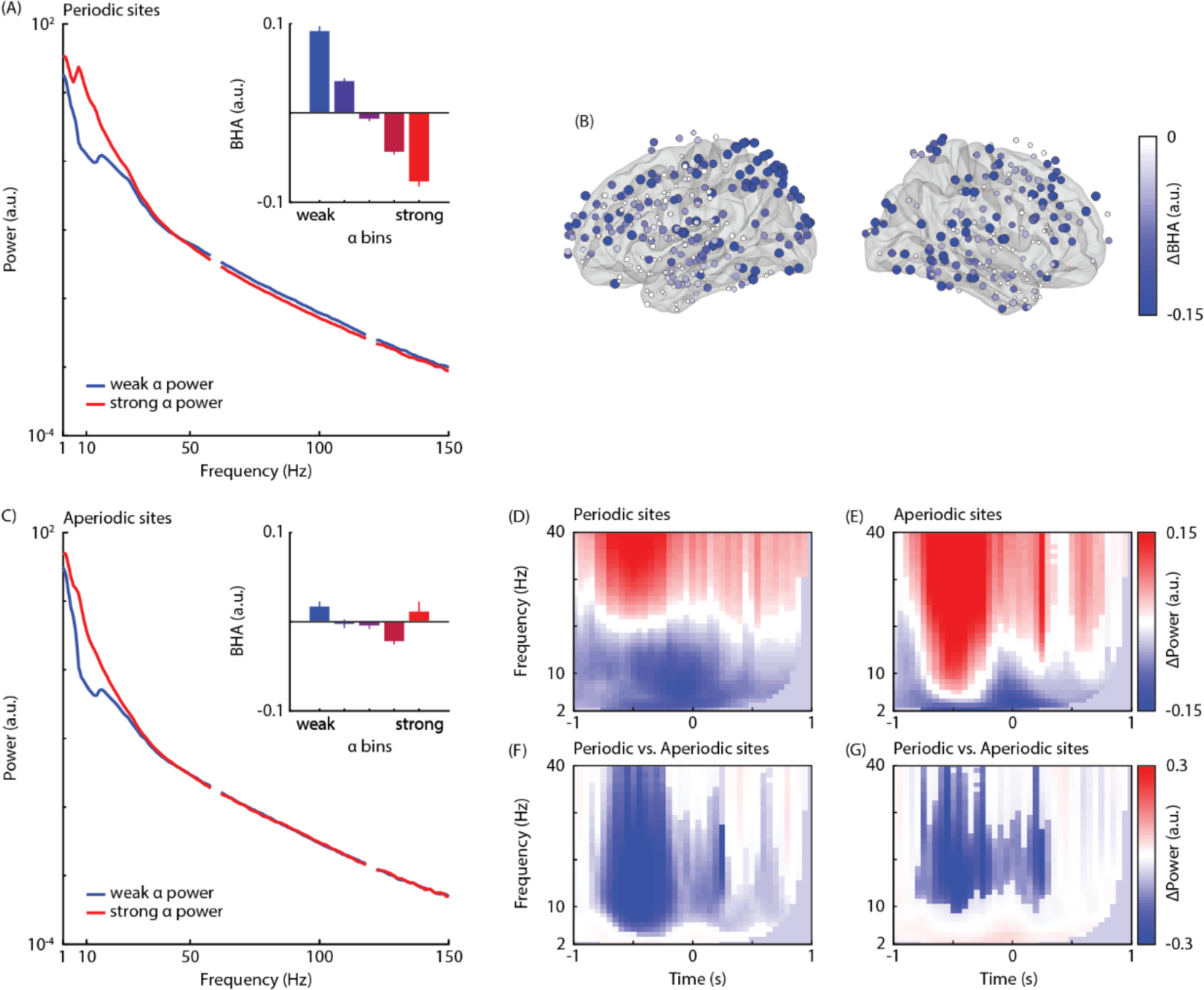
Correlation between prestimulus alpha power and prestimulus BHA. **A.** Averaged power spectrum computed during the 1-s prestimulus window in noise-locked epochs, shown separately for bins of strongest (red) and weakest (blue) prestimulus alpha power for periodic sites (spectra normalized with average power). The inset shows the averaged BHA (70–150 Hz) separately for five bins sorted from weakest (blue) to strongest (red) prestimulus alpha power (normalized with the average across bins) for periodic sites. BHA decreases with alpha power in periodic sites, consistent with functional inhibition. **B.** Anatomical map of the relationship between prestimulus alpha power and BHA power in periodic sites, estimated as the normalized difference in BHA between bins of strongest and weakest alpha power. The color and size of the dots are proportional to the magnitude of this difference. The map shows negative effects (BHA in bin 5 < BHA in bin 1). No significant positive effects were found. **C.** Same as (A) for aperiodic sites, respectively. BHA is unaffected by the aperiodic signal in the alpha frequency range. **D/E.** Time-frequency representations of the difference in low-frequency power between bins of strongest and weakest BHA estimated during the prestimulus window in noise-locked epochs (normalized by average power across the most extreme bins and masked to show significant effects only) in periodic and aperiodic sites, respectively. Negative (blue) and positive (red) clusters indicate that low-frequency power is weaker and stronger during states of strong BHA, respectively. **F/G**. Time-frequency representations of the comparison of the low-frequency power difference between prestimulus BHA bins across periodic and aperiodic sites (normalized by average power across sites and masked to show significant effects only). Panels F and G show the results for all sites and for a subset of periodic and aperiodic sites with matched high-frequency power difference, respectively. Negative clusters (blue) indicate that low-frequency power difference between BHA states is more negative in periodic sites. No positive clusters were found.

To determine whether this simultaneous correlation also exists during stimulus processing (i.e., window after the onset of the noise image), we sorted epochs based on alpha power estimated in the 1-s poststimulus window and computed simultaneous BHA. We found a significant interaction effect between periodic/aperiodic sites and poststimulus alpha bins on BHA (F(1.98, 2061.23)=44.01, p<0.001; bin 5 vs. bin 1 in periodic sites: t=−18.86, p<0.001; in aperiodic sites: t=−1.39, p=1), consistent with the results on the prestimulus window.

In sum, these results suggest that states of strong alpha oscillations—but not the aperiodic signal in the same frequency range—are linked to reduced BHA, consistent with functional inhibition. Importantly, this inhibitory effect was observed throughout the brain, and during moments of endogenous (i.e., prestimulus) as well as exogenous processing (i.e., poststimulus), suggesting that this is a general property of alpha oscillations.

### BHA negatively correlates with power in the alpha and neighboring bands

Is the inhibitory function specific to alpha oscillations? To address this question, we estimated time-frequency representations (TFRs) of low frequency power (2–40 Hz) during a 2-s time window centered on the onset of the noise image, separately for each electrode. Then, we sorted the epochs into five bins based on the single-trial estimates of BHA computed in the 1-s prestimulus window, and compared low frequency power between bins of strongest and weakest BHA. We carried out the group-level statistical analysis using cluster-permutation tests across sites to determine the frequency and time points of the difference in low-frequency power between the most extreme bins. If inhibition is specific to alpha, we expect that BHA sorting would be statistically unrelated to oscillatory power outside of the alpha range.

We ran separate tests for periodic and aperiodic sites. In periodic sites (Figure 2D), we found a significant negative cluster (t=−5165.32, p<0.001, from −1 to 0.9 s, 2–26 Hz peaking at 8 Hz), indicating that the strongest BHA bin was related to weaker low-frequency power, especially within the alpha band. In addition, we found a positive cluster (t=4736.05, p<0.001, from −1 to 0.95 s, 20–40 Hz), indicating stronger power in the strongest BHA bin. In aperiodic sites (Figure 2E), we found a significant negative cluster (t=−1933.54, p<0.001, from −1 to 0.85 s, 2–23 Hz peaking at 5 Hz), as well as two significant positive clusters (t=5365.82, p<0.001, from −0.9 to 0.35 s, 6– 40 Hz; t=603.89, p=0.005, from 0.5 to 0.75 s, 15–40 Hz).

To determine the effects specific to oscillatory activity, we subtracted the effect in aperiodic sites from the effect in periodic sites and tested for statistical significance using cluster permutation test. We found a significant negative cluster (t=−4787.24, p<0.001, from −0.85 to 0.8 s, 4–40 Hz peaking at 15 Hz; Figure 2F), indicating that, compared to aperiodic sites, states of strong BHA in periodic sites were associated with a greater power decrease in the alpha as well as theta, and beta bands. Note, however, that the difference in high-frequency power between states of strong and weak BHA was different between periodic (Figure supplement 2D) and aperiodic sites (Figure supplement 2E). Therefore, it is possible that the effects in low-frequency power between periodic and aperiodic sites (Figure 2F) may be driven by differences in high-frequency power (Figure supplement 2F). To address this issue, we replicated the analysis in a subset of periodic (N=90) and aperiodic (N=91) sites, with matched high-frequency power difference between states of strong and weak BHA (i.e., within ± 1/6 standard deviation from the median difference across all sites; Figure supplement 2G), and found that, compared to aperiodic sites, states of strongest BHA in periodic sites were associated with a greater power decrease in the alpha and beta band (t=−1411.55, p<0.001; from −0.8 to 0.3 s, 9–40 Hz peaking at 24 Hz; Figure 2G), consistent with the results for all sites. Additionally, the results of these analyses were replicated by sorting epochs based on frequency-normalized BHA estimates. Taken together, states of strong prestimulus BHA were related to lower prestimulus oscillations in the alpha+ band (alpha and neighboring frequencies), suggesting that the functional inhibition account is not exclusive to a narrowband alpha rhythm.

To corroborate these results, we sorted epochs into five bins based on prestimulus power in the beta band and then compared BHA across bins. We hypothesized a negative relationship between prestimulus beta power and BHA in sites with periodic beta activity (N=144 with mean peak at 20 Hz, SEM=0.26; Figure supplement 1) but not in sites with aperiodic activity (N=691). Similarly to the analysis of prestimulus alpha oscillations, we found a significant interaction effect between periodic/aperiodic sites and beta bins on BHA (F(1.87, 1558.68)=27.13, p<0.001; bin 5 vs. bin 1 in periodic sites: t=−6.73, p<0.001; Figure supplement 2A/B; in aperiodic sites: t=9.16, p<0.001; Figure supplement 2C). To rule out the possibility that these findings were driven by alpha-band harmonics, we replicated the analysis of BHA using a subsample of periodic sites (N=29) without alpha oscillations (beta-only periodic sites; Figure supplement 1A), and found a similar, although weaker, interaction effect (F(1.88, 1351.87)=3.15, p=0.036; bin 5 vs. bin 1 in periodic sites: t=−1.65, p=1; in aperiodic sites: t=9.29, p<0.001).

In sum, we found that states of strong prestimulus beta oscillations—but not the aperiodic signal in the same frequency range—are linked to reduced BHA, consistent with the results on alpha oscillations.

### Prestimulus alpha power negatively correlates with poststimulus BHA

Based on the functional inhibition account, we hypothesized that, by setting the state of neuronal excitability, prestimulus alpha oscillations affect the processing of incoming stimulus information. To test this hypothesis, we sorted epochs into five bins based on prestimulus alpha power and computed the BHA during sensory processing: i.e., in the 1-s poststimulus window. In noise-locked epochs, we found a significant main effect of alpha bin (F(3.50, 3641.44)=24.87, p<0.001), indicating that, across sites, poststimulus BHA (i.e., after noise image onset) (bin 5 vs. bin 1: t=−9.27, p<0.001) decreased with prestimulus alpha power (i.e., before noise image onset). Furthermore, we found a significant main effect of periodic/aperiodic sites (F(1, 1042)=67.67, p<0.001), with higher poststimulus BHA in periodic sites (t=8.23, p<0.001). Critically, we found a significant interaction effect between periodic/aperiodic sites and prestimulus alpha bins (F(3.562, 3711.934)=13.38, p<0.001), indicating that, in periodic sites, BHA during the processing of the noise image decreased with alpha power estimated during the window preceding the noise image (bin 5 vs. bin 1: t=−11.48, p<.001; Figure 3A). By contrast, in aperiodic sites, poststimulus BHA was unrelated to prestimulus alpha power (bin 5 vs. bin 1: t=−1.66, p =0.880). We obtained independent evidence for this effect when analyzing syllable-locked epochs. Specifically, we found a significant interaction effect between periodic/aperiodic sites and prestimulus alpha bins (i.e., before syllable onset) on poststimulus BHA (i.e., after syllable onset) (F(3.56, 3711.93)=13.38, p<0.001, bin 5 vs. bin 1 in periodic sites: t=−5.15, p<0.001; Figure 3B; in aperiodic sites: t=−0.89, p=1), suggesting that this effect generalizes across visual and auditory sensory stimulation.

**Figure 3.**
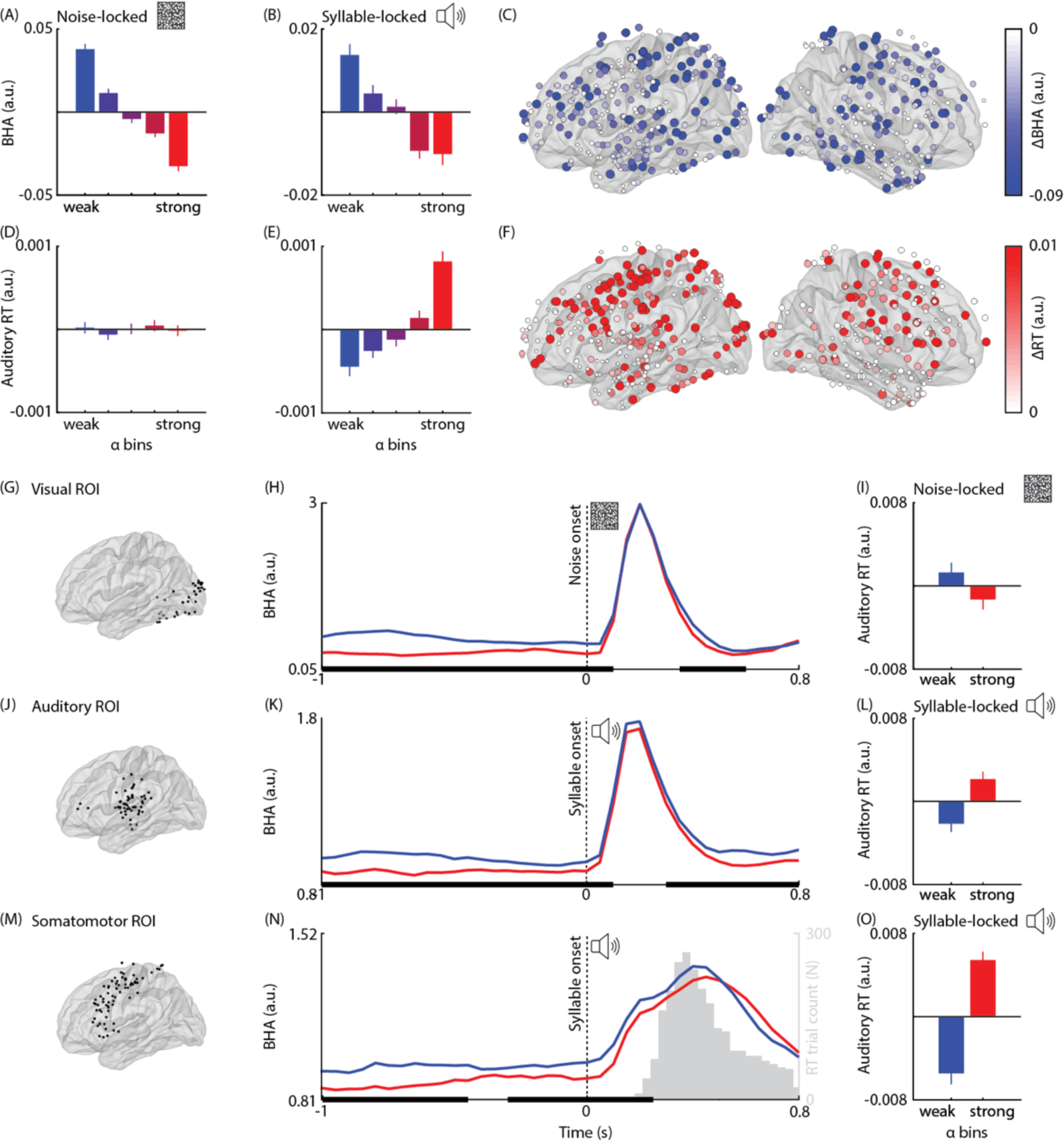
Correlation between prestimulus alpha power, poststimulus BHA, and reaction times. **A/B.** Averaged poststimulus BHA, shown separately for five bins sorted from weakest (blue) to strongest (red) prestimulus alpha power in periodic sites in noise-locked and syllable-locked epochs, respectively. In periodic sites, BHA after the onset of the noise image and the syllable decreases with prestimulus alpha power, consistent with functional inhibition. **C**. Anatomical map of the relationship between prestimulus alpha power and poststimulus BHA in periodic sites, estimated as the normalized difference between bins of strongest and weakest alpha power. The color and size of the dots are proportional to the experimental effect. The map shows negative effects (BHA in bin 5 < BHA in bin 1) in noise-locked epochs. No significant positive effects were found. **D/E.** Same as in (A) and (B) for RT in noise- and syllable-locked epochs, respectively. RT increases with prestimulus alpha power in syllable-locked epochs, but not in noise-locked epochs. **F**. Same as in (C) for RT. The map shows positive effects (RT in bin 5 > RT in bin 1) in syllable-locked epochs. No significant negative effects were found. **G.** Anatomical map of the visual ROI, including periodic sites that were functionally and anatomically related to visual cues. **H.** BHA time-course shown separately for bins of weakest (blue) and strongest (red) prestimulus alpha power for noise-locked epochs in the visual ROI. Bold horizontal black lines indicate significant differences using cluster permutation testing. States of strong prestimulus alpha power are related to a BHA reduction during the prestimulus and poststimulus windows, consistent with functional inhibition. **I.** Averaged RT, shown separately for the weakest (blue) and strongest (red) bin of prestimulus alpha power in noise-locked epochs in the visual ROI. RT is affected by alpha power before the syllable in both auditory and somatomotor areas, but not before the noise image in visual areas. **J.** Same as (G) for the auditory ROI, including periodic sites that were functionally and anatomically related to target syllables. **K/L.** Same as (H), and (I) for syllable-locked epochs in the auditory ROI. **M**. Same as (G) for the somatomotor ROI, including periodic sites that were functionally and anatomically related to motor responses. **N/O.** Same as (H), and (I) for syllable-locked epochs in the somatomotor ROI.

We also compared BHA time-series between bins of strongest and weakest prestimulus alpha power separately for three regions of interest (ROI) based on functional and anatomical localizers. In the visual ROI (N=37; Figure 3G), we found two significant negative clusters, indicating that states of weak prestimulus alpha power (bin 1) coincided with weaker prestimulus BHA (t=−116.59, p<0.001; from −1 to 0.1 s) and were followed by weaker poststimulus BHA (t=−24.52, p=0.013, from 0.35 to 0.6 s) in noise-locked epochs (Figure 3H). These results were replicated during scene- (Figure 3A supplement) and face-locked epochs (Figure 3C supplement). In the auditory ROI (N=55; Figure 3J), we found two significant negative clusters, indicating that states of weak prestimulus alpha power (bin 1) coincided with weaker prestimulus BHA (t=−84.75, p<0.001, from −1 to 0.1 s) and were followed by weaker poststimulus BHA (t=−37.89, p<0.001, from 0.3 to 0.85 s) in syllable-locked epochs (Figure 3K). In the somatomotor ROI (N=64; Figure 3M), we found two significant clusters, indicating that states of weak prestimulus alpha power (bin 1) coincided with weaker prestimulus BHA (t=−59.07, p=0.002; from −1 to −0.45 s) and were followed by weaker poststimulus BHA (t=−46.61, p=0.002, from −0.3 to 0.25 s) in syllable-locked epochs (Figure 3N). The ROI results suggest that this effect generalizes across visual, auditory and somatomotor areas.

Taken together, these results show that states of strong prestimulus alpha oscillations are followed by reduced poststimulus BHA across sensory modalities and brain areas, consistent with functional inhibition.

### Prestimulus alpha power positively correlates with reaction times

Based on the functional inhibition account, we hypothesized that states of strong prestimulus alpha oscillations in task-relevant brain regions are followed by behavioral changes (e.g., slower RTs). To test this, we compared the RTs of the auditory discrimination task across states of weak and strong prestimulus alpha power.

In syllable-locked epochs, we found a significant main effect of prestimulus alpha bins on RTs (F(3.39, 3534.06) t=62.60, p<0.001), indicating that RTs increased with prestimulus alpha power in both periodic (bin 5 vs. bin 1: t=11.94, p<0.001; Figure 3E) and aperiodic sites (bin 5 vs. bin 1: t=12.92, p<0.001). The mean RT difference between states of strong and weak prestimulus alpha power in periodic sites was 0.011 s (SEM= 27) with a maximum of 0.235 s in the caudal division of the middle frontal gyrus, while in aperiodic sites it was 0.018 s (SEM= 23) with a maximum of 0.143 s in the postcentral gyrus. An anatomical map of the RT difference in periodic sites revealed that slow RTs were preceded by strong pre-syllable alpha power across the brain (Figure 3F). In addition, we found a significant interaction effect (F(3.39, 3534.06)=3.09, p =0.021), suggesting a trend for slower RTs in the strongest alpha bin in aperiodic sites (periodic vs. aperiodic sites for bin 5: t=−2.95, p=0.067). Note that the effect of prestimulus alpha power on RTs was present during syllable-locked epochs in both auditory (bin 5 vs. bin 1: t(54)=3.32, p=0.002) and somatomotor ROIs (bin 5 vs. bin 1: t(63)=6.94, p<0.001). By contrast, no significant effects were found in noise- (Figure 3I), scene- (Figure 3B supplement) and face-locked epochs (Figure 3D supplement) in the visual ROI (p>0.05), indicating that auditory RTs were affected only by alpha power right before the target auditory stimulus.

In sum, these results demonstrate that states of strong prestimulus alpha oscillations—as well as the aperiodic signal in the same frequency range—are followed by slower RTs. However, it should be noted that this result alone does not constitute evidence for the functional inhibition account of alpha oscillations since it is not specific to oscillatory activity.

### The correlation between prestimulus alpha power and RT is mediated by poststimulus BHA

Based on the functional inhibition account, we hypothesized that the RT effect associated with prestimulus alpha oscillations (rather than the aperiodic signal) was specifically mediated by a modulation of poststimulus BHA. We estimated the mediation effect by analyzing the trial-by-trial interrelation between alpha power, RTs, and BHA using a causal step approach based on generalized linear model regression (GLM; Judd & Kenny, 1981; Figure 4A/C, see Methods). Confirming the results of the binning analysis (Figure 3B), we found that prestimulus alpha was negatively correlated with poststimulus BHA in periodic sites (t(524)=−3.569, p<0.001; a<0 in Figure 4A), consistent with functional inhibition, whereas it was positively correlated with poststimulus BHA in aperiodic sites (t(518)=2.22, p=0.027) (a>0 in Figure 4B), consistent with a change in the offset of the power spectrum (i.e., upward shift at all frequencies). In addition, we found that poststimulus BHA was negatively correlated with RT across all sites (periodic: t(524)=−2.03, p=0.043; aperiodic: t(518)=−5.85, p<0.001) (b>0 in Figure 4B), even after controlling for prestimulus alpha power (periodic: t(524)=−1.98, p=0.048; aperiodic: t(518)=−5.92, p<0.001) (b’<0 in Figure 4B). Moreover, we found that prestimulus alpha was positively correlated with RT in periodic (T(524)=7.33, p<0.001) and aperiodic sites (T(518)=11.18, p<0.001) (c in Figure 4B), even after controlling for poststimulus BHA (periodic: t(524)=6.77, p<0.001; aperiodic: t(518)=10.93, p<0.001; c’>0 in Figure 4B), suggesting partial mediation (see Methods).

**Figure 4.**
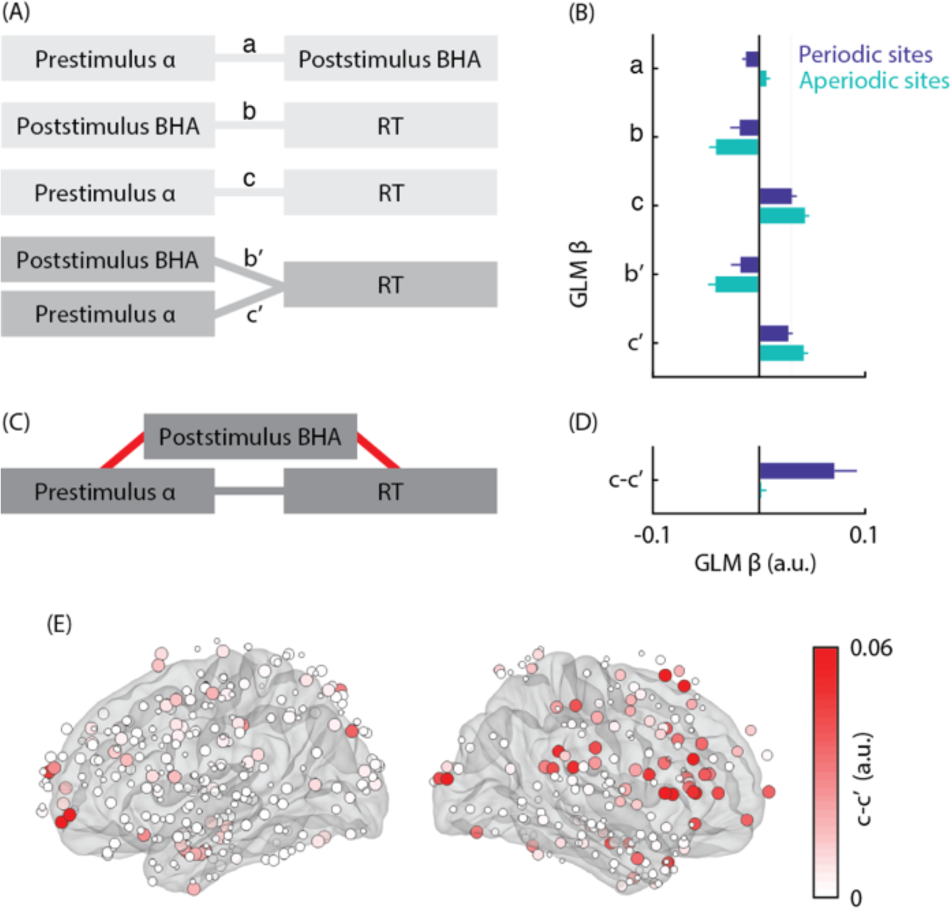
Mediation between prestimulus alpha power, poststimulus BHA, and reaction time. **A.** Schematic illustration of the causal path mediation analysis using four GLMs, characterizing the interrelation between prestimulus alpha oscillations (i.e., the independent variable), RT (i.e., the dependent variable), and poststimulus BHA (i.e., the mediator) in syllable-locked epochs: *a* represents the relationship of independent variable to the mediator, *b* the mediator to dependent variable, *c* the independent variable to dependent variable, *b’* the mediator to dependent variable adjusted for independent variable, and *c’* the independent variable to dependent variable adjusted for mediator. The variables and the relationships among them are represented by rectangles and lines, respectively. **B**. Averaged effects of the causal path mediation analysis showing the interrelation between prestimulus alpha power, poststimulus BHA, and RT, shown separately for periodic (blue) and aperiodic sites (green). These effects are estimated using the averaged zero-order (a–c) and partial GLM coefficients (b’/c’) and their comparison (c-c’) based on causal path mediation analysis in (A). **C.** Theoretical mediation model in which the independent variable leads to the dependent variable via an indirect effect through the mediator (red path). **D.** Averaged indirect effect reflecting the mediation between prestimulus alpha power and RT via poststimulus BHA, shown separately for periodic (blue) and aperiodic sites (green). In periodic sites, the effect of prestimulus alpha power on RT is significantly reduced after accounting for poststimulus BHA (c-c’>0), demonstrating an indirect effect, consistent with functional inhibition. In aperiodic sites, the relationship between prestimulus alpha power and RTs is unaffected by poststimulus BHA (c-c‘=0), suggesting no indirect effect. **E**. Anatomical map of the magnitude of the indirect effect in periodic sites. The color and size of the dots are proportional to the experimental effect.

To test for mediation, we estimated the indirect effect by computing the reduction in the effect of prestimulus alpha power on RT after accounting for poststimulus BHA. We found a significant indirect effect in periodic sites (t(524)=3.41, p<0.001), supporting mediation, but not in aperiodic sites (t(518)=0.26, p=0.795; c-c’ in Figure 4D). This effect was indeed larger in sites with periodic activity (unpaired t(1042)=3.22, p=0.001). An anatomical map of the indirect effect in periodic sites revealed this effect occurred across several sites with maxima in the pars triangularis and the rostral division of the middle frontal gyrus (Figure 4E).

Together, these findings support the functional inhibition account by demonstrating that the modulation of poststimulus excitability mediates the behavioral effects of alpha oscillations, rather than the aperiodic signal in the same frequency range.

### Prestimulus alpha power negatively correlates with neural stimulus feature encoding

Based on the functional inhibition account, we hypothesized that the effect of prestimulus alpha power on poststimulus neuronal excitability may affect how stimulus features are encoded in BHA estimates. Specifically, we tested whether prestimulus alpha power affects (1) decoding *accuracy*, which reflects sensory precision or (2) decoding *confidence*, which reflects the strength of sensory encoding. To test this, we used “spatial decoding” (Gwilliams & King, 2020): separately for each electrode, we predicted syllable identity (labels: /ga/ vs. /pa/) by fitting logistic regression decoders on the single-trial temporal BHA patterns as input (see Figure 5 supplement for a comparison with the low-passed signal). First, decoder accuracy was estimated as the similarity between the probabilistic prediction (normalized distance from the hyperplane) and true labels using the area under the curve (AUC). Second, stimulus encoding strength or decoder confidence was estimated as the maximum probabilistic prediction, regardless of its accuracy (i.e., whether or not it matched the true label). Across sites there was a significant positive correlation between decoder accuracy and confidence averaged over trials (Spearman rho=0.15, p<0.001), demonstrating that sites which encode syllable identity more reliably also tend to exhibit higher confidence in those predictions. Moreover, to understand how these decoder metrics were related to the participant’s behavior on a trial-by-trial basis, we correlated RTs with single-trial accuracy (binarized into correct and incorrect) and confidence of the decoder predictions using GLM for each site. Across sites there was a significant negative correlation between RTs and both decoder accuracy (t(1043)=−2.75, p=0.006) and confidence (t(1043)= −3.45, p<0.001), indicating that higher decoder accuracy and confidence were related to faster RTs.

**Figure 5.**
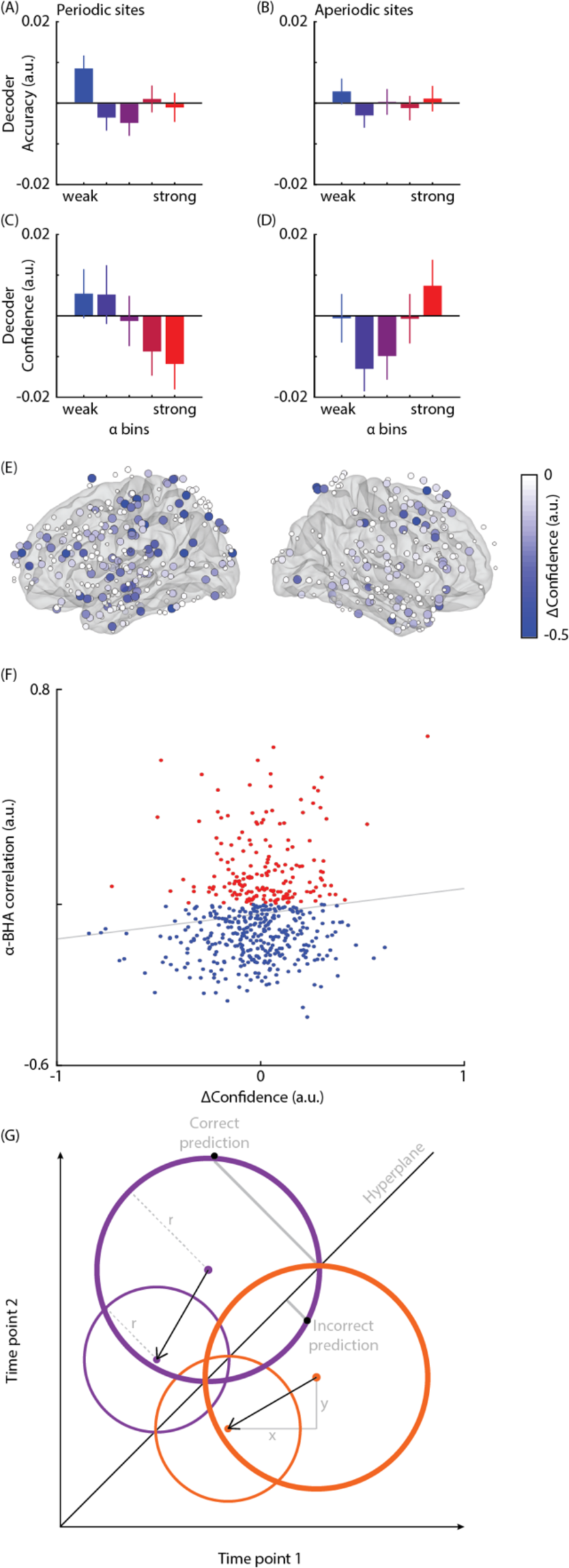
Correlation between prestimulus alpha power and neural stimulus decoding. **A**. Averaged normalized decoder accuracy (AUC) for decoding syllable identity from BHA shown separately for five bins sorted from weakest (blue) to strongest (red) prestimulus alpha power in periodic sites. **B.** Same as (A) in aperiodic sites. Decoder accuracy is affected by prestimulus alpha power in neither periodic nor aperiodic sites. **C.** Same as (A) for decoder confidence. **D.** Same as (C) for aperiodic sites. Decoder confidence decreases with prestimulus alpha power in periodic sites whereas it increases in aperiodic sites. **E.** Anatomical map of the magnitude of the effect of the decoder confidence in periodic sites. The color and size of the dots are proportional to the experimental effect. The map shows negative effects (confidence in bin 5 < confidence in bin 1). No significant positive effects were found. **F.** Across-site correlation between the effect of prestimulus alpha power on decoder confidence and the relationship between ongoing alpha power and BHA (i.e., index of functional inhibition). The dots represent the magnitude of the functional inhibition index (on y axis) and the effect of prestimulus alpha power on confidence (on x axis) for each electrode. Sites that show a negative relationship between ongoing alpha power and BHA, reflecting functional inhibition, are represented in blue. The grey line indicates the least-squares fit. A positive correlation indicates that, during states of strong prestimulus alpha power, sites with stronger alpha-related BHA suppression also show lower decoder confidence. **G.** Model of the relationship between alpha oscillations and decoding performance. In a two-dimensional activation space, each trial is characterized by the activity values estimated at two time points from a single electrode (i.e., activation point). Response distributions are shown by purple (for stimulus A) and orange circles (for stimulus B) with center and radius representing response mean and variability, respectively. Learning a linear classifier (A vs B) is equivalent to learning the hyperplane (diagonal line) that best separates the two distributions: if distance between an activation point and the hyperplane > 0, then the classifier predicts “A”, otherwise “B”. This model assumes that states of low excitability indexed by strong alpha oscillations are related to reduced distributions’ mean (arrow), and variability, resulting in a decrease of the absolute distance of the activation points from the hyperplane (i.e., lower decoder confidence), while leaving the distributions’ overlap (i.e., decoder accuracy) unaffected.

Next, we compared decoder accuracy (AUC) across alpha bins in periodic and aperiodic sites and found no significant effects (p>0.05; Figure 5A/B), suggesting that the precision of neural stimulus representation/encoding was unlikely affected by prestimulus alpha power. We also compared decoder confidence across alpha bins in periodic and aperiodic sites and found a significant interaction effect (F(3.96, 4125.61)=2.59, p=0.036), such that stimulus encoding strength nominally decreased with alpha power in periodic sites (bin 5 vs. bin 1: t=−1.89, p=1; Figure 5C), while it nominally increased in aperiodic sites (bin 5 vs. bin 1: t=0.87, p=1; Figure 5D), although these post-hoc effects were not significant after correcting for multiple comparisons. Additionally, this interaction effect was significant in trials with correct predictions (F(4.00, 4166.49)=3.54, p=0.007) and trending toward significance in trials with incorrect predictions (F(3.99, 4148.44)=2.06, p=0.084), suggesting that strong prestimulus alpha power in periodic sites decreased stimulus encoding strength (and vice versa in aperiodic sites) regardless of the accuracy of stimulus encoding. An anatomical map of the effect on confidence in periodic sites revealed a widespread reduction following states of strong prestimulus alpha power across the brain (Figure 5E).

We also tested whether, in periodic sites, the effects of alpha power on decoder accuracy and confidence (i.e., bin 5 vs. bin 1) were dependent on two site-specific characteristics: the negative relationship between ongoing alpha power and BHA (i.e., index of functional inhibition), and overall decoder accuracy. We found no across-site correlation (p>0.05) between overall decoder accuracy and the effects of alpha power on decoder accuracy and confidence, or between the alpha-BHA relationship and the effect on decoder accuracy, indicating that decoder accuracy was unrelated to the confidence effect and to prestimulus power. By contrast, we found a significant positive across-site correlation between the alpha-BHA relationship and the effect of alpha power on decoder confidence (Spearman rho=0.103, p=0.019; Figure 5F), suggesting that, during states of strong prestimulus alpha power, the stronger the inhibition associated with alpha power, the lower the decoder confidence. We corroborated this finding in sites with a negative alpha-BHA relationship, reflecting functional inhibition: we found a significant effect of alpha bins on decoder confidence (F(3.891, 1338.467)=3.69, p=0.006), indicating that decoder confidence decreased with prestimulus alpha power (bin 5 vs. bin 1: t=−2.86, p=0.039).

In sum, these findings extend the functional inhibition account by demonstrating that states of strong prestimulus alpha power reduce the strength, but not the precision, of neural stimulus representations.

### Baseline shift of alpha oscillations affects ERPs but not BHA

Finally, we assessed if the observed relationship between alpha oscillations and BHA is determined by mechanisms other than functional inhibition (i.e., baseline shift of non-zero-mean oscillations) that affect event-related potentials (ERP; Nikulin et al., 2007; Mazaheri & Jensen, 2008; Iemi et al., 2019). Baseline shift predicts that the relationship between alpha power and neural signals depends on the direction of the baseline shift (which can also be viewed as polarity) of non-zero-mean oscillations (Figure 6A/B), whereas functional inhibition predicts a negative relationship between alpha power and neural signals (reflecting excitability) regardless of oscillatory polarity. To test for these mechanisms, we identified 323 sites with a negative oscillatory mean (Figure 6C/E) and 202 sites with a positive oscillatory mean (Figure 6D/F) using the baseline shift index (BSI), and analyzed how prestimulus oscillations with different polarities affect ERP and BHA.

**Figure 6.**
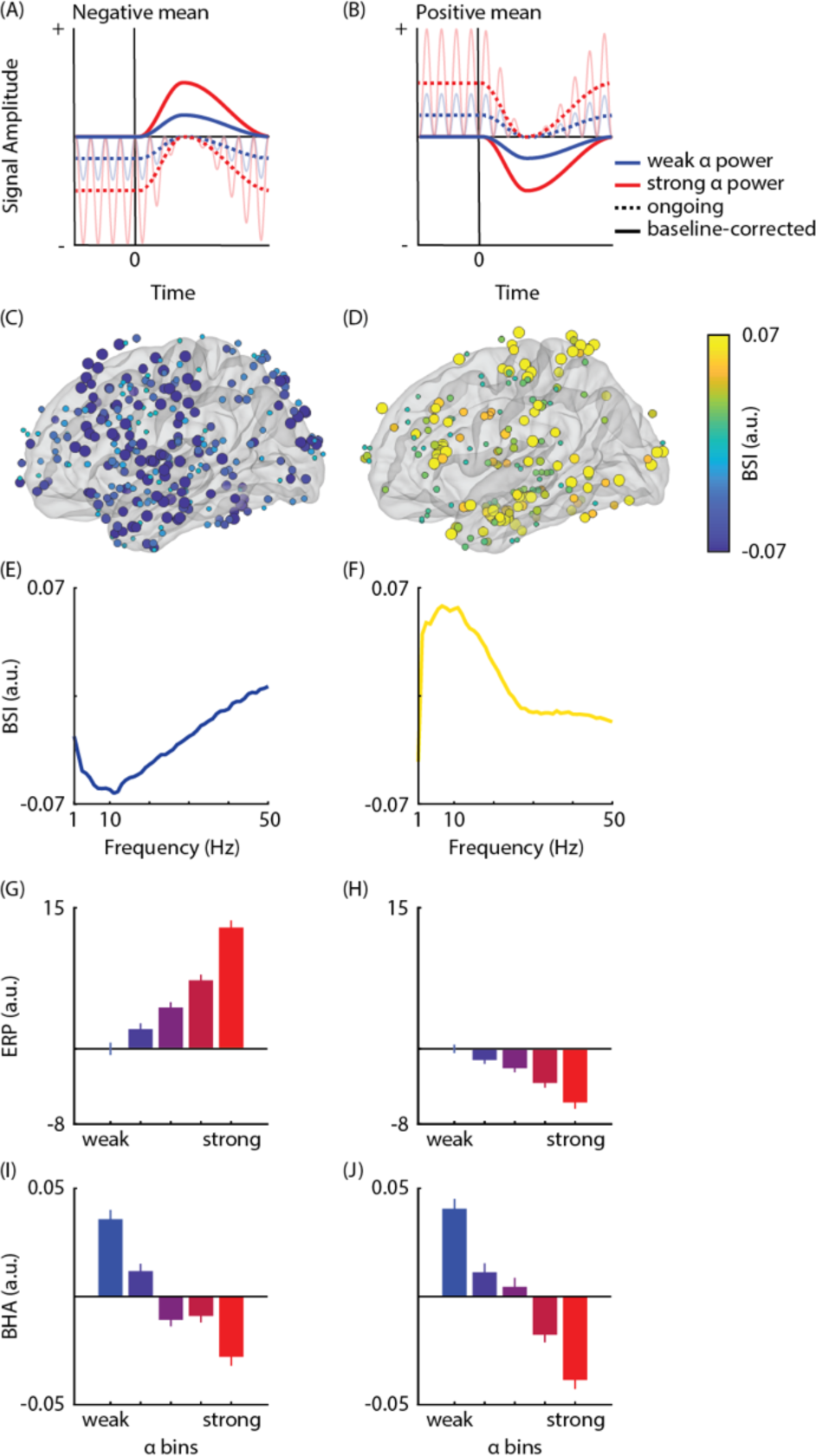
Correlation between alpha power and prestimulus BHA accounting for baseline shift of non-zero-mean oscillations. **A/B.** Neural oscillations are modelled as sinusoidal waveforms (opaque lines) varying asymmetrically around a non-zero mean. Unlike zero-mean oscillation, trial averaging of non-zero-mean oscillations does not eliminate non-phase-locked oscillations in the prestimulus window, resulting in an ERP prestimulus baseline with an offset (dotted lines). During event-related desynchronization (ERD) by stimulus onset (vertical line), the averaged signal gradually approaches the zero line of the signal. When the poststimulus signal is corrected with the prestimulus non-zero baseline, a slow shift of the ERP signal appears (thick lines), mirroring the ERD. The baseline-shift mechanism predicts that: (1) the polarity of the non-zero mean baseline determines the directionality of the effect of prestimulus oscillations on the ERP signal (in raw values); (2) the poststimulus ERP signal is amplified (in absolute values) during states of strong prestimulus alpha power. **C/D.** Anatomical map of the magnitude of the baseline-shift-index (BSI) averaged across the alpha band shown separately for periodic sites with negative- and positive-mean alpha oscillations, respectively. **E/F.** Magnitude of the baseline-shift-index (BSI) represented for frequencies between 1 and 50 Hz, separately for sites with negative- and positive-mean alpha oscillations, respectively. **G/H.** ERP magnitude averaged across the 1-s poststimulus window, shown separately for five bins sorted from weakest (blue) to strongest (red) prestimulus alpha power in periodic sites with negative and positive mean oscillations, respectively. The estimates were normalized, first, with the average across bins and, second, with the magnitude of the weakest bin, which is assumed to best reflect the zero line of the signal. The relationship between prestimulus alpha power and poststimulus ERP (in raw values) depends on the non-zero-mean property of alpha oscillations. States of strong prestimulus alpha power results in ERP amplification (in absolute values), consistent with baseline shift. **I/J.** Averaged poststimulus BHA shown separately for five bins sorted from weakest (blue) to strongest (red) prestimulus alpha power in periodic sites with negative and positive mean oscillations, respectively (normalized with the average across bins). There is a negative relationship between prestimulus alpha power and poststimulus BHA regardless of the polarity of the oscillatory mean, consistent with functional inhibition.

We found a significant interaction effect between polarity (i.e., sites with positive or negative mean) and prestimulus alpha bin on the prestimulus (F(1.79, 935.91)=120.45, p<0.001) and poststimulus ERP (F(1.67, 873.44)=139.55, p<0.001; Figure 6G/H). Compared to the weakest alpha bin, the prestimulus ERP baseline of the strongest alpha bin was characterized by a more negative voltage in sites with a negative mean (bin 5 vs. bin 1: t=−19.48, p<0.001), and more positive voltage in sites with a positive mean (bin 5 vs. bin 1: t=10.09, p<0.001). Furthermore, compared to the weakest alpha bin, the ERP averaged over the poststimulus window of the strongest alpha bin was characterized by a more positive voltage shift in sites with a negative mean (bin 5 vs. bin 1: t=23.99, p<0.001; Figure 6G), and more negative voltage shift in sites with a positive mean (bin 5 vs. bin 1: t=−8.39, p<0.001; Figure 6H). In other words, states of strong prestimulus alpha power are related to a stronger non-zero ERP baseline, which is followed by an ERP amplification in the poststimulus window, consistent with a baseline shift of non-zero-mean alpha oscillations.

By contrast, we found a significant main effect of prestimulus alpha bins on both prestimulus (F(2.11, 1104.11)=99.79, p<0.001) and poststimulus BHA (F(3.51, 1834.91)=28.93, p<0.001), indicating that BHA decreased with prestimulus alpha power in sites with both a negative (bin 5 vs. bin 1 for prestimulus BHA: t=−12.78, p<0.001; poststimulus BHA: t=−7.85, p<0.001; Figure 6I), and positive mean (prestimulus BHA: t=−12.86, p<0.001; poststimulus BHA: t=−6.40, p<0.001; Figure 6L), consistent with functional inhibition. Indeed, there was no significant interaction effect between polarity and prestimulus alpha bin on prestimulus or poststimulus BHA (p>0.05). In addition, we found a significant main effect of polarity on both prestimulus (F(1, 523)=8.48, p=0.004) and poststimulus BHA (F(1, 523)=7.30,p=0.007), indicating that sites with a negative oscillatory mean were characterized by overall greater BHA (prestimulus BHA: t=2.91, p=0.004; poststimulus BHA: t=2.70, p=0.007). We replicated these effects in a control analysis using the Amplitude Fluctuation Asymmetry Index (AFAI: Mazaheri & Jensen, 2008) to identify sites with a negative and positive asymmetries in oscillatory peaks (Figure 6 supplement). In sum, these results demonstrate that the negative relationship between alpha power and BHA is consistent with functional inhibition, rather than baseline shift of non-zero mean alpha oscillations.

## Discussion

### Simultaneous relationship between alpha oscillations and excitability

We hypothesized that alpha oscillations reflect functional inhibition, regulating the excitability state of the neural system. To test this, we analyzed the relationship between alpha power and BHA (often referred to as “high gamma”) which is considered a proxy for neuronal ensemble excitability. Initially thought to reflect net local neuronal firing—i.e., multiunit activity (Manning et al., 2009; Nir et al., 2007; Ray et al., 2008; Ray & Maunsell, 2011; Rich & Wallis, 2017; Whittingstall & Logothetis, 2009), it was shown more recently that BHA mainly indexes calcium-dependent dendritic processes that, albeit correlated with firing probability, are separable from it (Leszczyński et al., 2020). This suggests that in relative terms, BHA is a more direct measure of neuronal excitability. There is also indication that BHA signals may volume-conduct over greater distances than multiunit activity signals (Leszczyński et al., 2020), making BHA a more useful measure of neuronal ensemble excitability. Our findings confirmed the functional inhibition account by showing that states of strong ongoing oscillations in the alpha+ band (i.e., alpha and neighboring frequencies) were related to reduced BHA, indicating a state of low neuronal excitability.

These findings corroborate studies reporting that strong alpha oscillations are related to reduced neuronal firing in animal models (Haegens et al., 2011; Watson et al., 2018) and reduced fMRI BOLD signal in humans (Chapeton et al., 2019; Mayhew et al., 2013). Additionally, they are also consistent with numerous studies suggesting a negative relationship between low-frequency power, including the alpha band, and high-frequency power, including BHA, during sensory, motor and task processing. Specifically, increased high-frequency power co-occured with decreased low-frequency power after the presentation of visual (Lachaux et al., 2005; Martin et al., 2019; Nir et al., 2007, 2017; Rodriguez et al., 2004; Fisch et al., 2009; Miller et al., 2014; Podvalny et al., 2015; Fries et al., 2001; Scheeringa et al., 2011; Hwang & Andersen, 2011; Rickert et al., 2005; Lundqvist et al., 2020), auditory (de Pesters et al., 2016; Potes et al., 2014), and somatosensory stimuli (Fontolan et al., 2014), and during motor responses (Crone et al., 1998; de Pesters et al., 2016; Jiang et al., 2020; Miller et al., 2007) and task/cognitive processing (Asher et al., 2007; Hwang & Andersen, 2011). Several M/EEG studies observed a negative relationship between low-frequency power decrease and, what is reported as, narrowband gamma increase (Bauer et al., 2006; Kloosterman et al., 2019; van Ede et al., 2014; Wyart & Tallon-Baudry, 2009); though it remains unclear whether this high-frequency signal reflects genuine oscillatory activity or broadband power. Importantly, the negative relationship between low- and high-frequency power was recently linked to changes in cognitive function (Nir et al., 2007; Proskovec et al., 2019; Tran et al., 2020; Voytek et al., 2015).

In this study, rather than analyzing how BHA and alpha power change as a function of sensory/task processing, we systematically estimated the trial-by-trial correlation between these signals. We analyzed, first, a wide range of brain areas spanning from primary sensory cortices to frontal regions, and second, time windows of endogenous (prestimulus) and exogenous (poststimulus) processing. In this way, we overcome some limitations in previous studies estimating neural signals in a limited set of brain regions (e.g., Barne et al., 2020; Haegens et al., 2011), and/or following experimental events (e.g., saccades: Barczak et al., 2019), which cause phase-reset and thereby impact ongoing oscillatory estimates. Our findings show a negative relationship between alpha power and BHA occurring across the whole brain and both during the prestimulus and poststimulus window, consistent with the idea that functional inhibition reflects a general property of alpha oscillations.

Third, some investigators have questioned whether the negative correlation between low- and high-frequency power reflects an oscillatory modulation driven by the alpha rhythm or whether it is due to a change of the aperiodic signal (McNair et al., 2019; Podvalny et al., 2015; Voytek et al., 2015), since the power spectrum contains not only oscillatory/periodic activity, but also an aperiodic signal (1/f background noise), parametrized by an offset and a slope (Donoghue et al., 2020). While an increase in the offset of the aperiodic signal may boost power at all frequencies, an increase in aperiodic slope may manifest as a simultaneous increase in low-frequency power and a decrease in high-frequency power. Critically, sorting trials by the power in a predefined frequency band (e.g., alpha) in different bins has been shown to affect both the slope and offset of the aperiodic signal (Iemi et al., 2019). Therefore, it is possible that the reduced BHA during states of strong alpha power can be explained by a steeper slope of the aperiodic signal. Moreover, the aperiodic signal is thought to reflect a physiological function (Voytek et al., 2015) that is, at least partially, independent from the periodic signal. Accordingly, distinguishing between an oscillatory or aperiodic modulation driving the alpha-BHA relationship is critical for understanding the underlying neural mechanisms. In periodic sites, containing both periodic/oscillatory and aperiodic activity, it is not possible to distinguish between the oscillatory and aperiodic modulations because we expect the same negative relationship between alpha power and BHA, regardless of the underlying modulation. By contrast, in aperiodic sites, we can make separate predictions for the oscillatory and aperiodic modulations. If, on the one hand, the alpha-BHA relationship reflects an aperiodic modulation (i.e., change of the aperiodic slope), then we expect a negative relationship in aperiodic sites: namely, states of strong aperiodic power in the alpha range may be related to a steeper aperiodic slope, resulting in lower BHA. If, on the other hand, the alpha-BHA relationship reflects an oscillatory modulation, then we expect a positive relationship in aperiodic sites: namely, states of strong aperiodic power in the alpha range may be related to a higher offset of the power spectrum (i.e., upward shift at all frequencies), resulting in higher BHA. Our results are consistent with an oscillatory modulation, showing a negative relationship between BHA and low-frequency power in periodic sites, and a positive relationship in aperiodic sites (alpha: Figure 4B; beta: Figure supplement 2A/C). It should be noted that the distinction between periodic and aperiodic sites was based on the peak detection in the trial-averaged prestimulus power spectrum because it provides a more accurate spectral estimate than individual trials. In fact, due to low signal-to-noise ratio, it is difficult to distinguish whether a peak detected in a single trial reflects spurious or genuine oscillatory activity. Note that, even if some trials in aperiodic sites contain genuine peaks, this would work against the hypothesis of oscillatory modulation.

Fourth, we analyzed the relationship between BHA and power in a range of low frequencies (2– 40 Hz). Thus, we overcome a limitation of some previous studies which only focused on a predefined frequency-band (e.g., alpha: Potes et al., 2014). We found that strong BHA occurred during states of weak power in the alpha+ band (i.e., alpha and neighboring frequencies including beta and theta), suggesting that the functional inhibition account is not exclusive to a narrowband alpha rhythm. Additionally, we found that BHA was decreased in states of strong beta power in sites with periodic activity, consistent with numerous reports of a negative correlation between beta and high-frequency power (Bauer et al., 2006; Fisch et al., 2009; Fontolan et al., 2014; Kloosterman et al., 2019; Watson et al., 2018). While it is possible that frequencies around the alpha band may reflect a similar function, it is important to highlight that the relationship between ongoing power and BHA was more prominent and sustained in time within the alpha band (Figure 2). Additionally, it is possible that analysis techniques such as frequency smoothing may have contributed to the spreading of the effect beyond the alpha band. Finally, we found that ongoing delta-band power (<4 Hz) was unrelated to BHA (Figure 2F/G), suggesting that the relationship between BHA and low-frequency power was unlikely determined by a change of the slope of the aperiodic signal affecting all low frequencies (including delta). This finding is particularly important as it further demonstrates that the relationship between alpha oscillations and BHA reflects a true oscillatory modulation.

In sum, these findings confirm the functional inhibition account and extend previous work by establishing the anatomical, temporal, and spectral characteristics of the relationship between alpha oscillations and neuronal excitability.

### Relationship between prestimulus alpha power and poststimulus excitability

Based on the functional inhibition account, we hypothesized that states of strong prestimulus alpha oscillations are followed by low neuronal excitability during stimulus processing (i.e., in the poststimulus window). We found that states of strong alpha power were followed by a reduction in BHA during both the processing of visual and auditory stimuli, as well as across sensory and non-sensory regions, suggesting that this phenomenon reflects a general property of alpha oscillations. These results are consistent with functional inhibition and with previous studies showing that ongoing alpha power is negatively correlated with the BOLD signal in sensory and non-sensory areas (Becker et al., 2008; Scheeringa et al., 2011; Walz et al., 2015) and early ERP components (Baslar & Stampfer, 1985; Becker et al., 2008; Iemi et al., 2019; Jasiukaitis & Hakerem, 1988; Rahn & Başar, 1993; Roberts et al., 2014). It is important to note the inhibitory effect of alpha oscillations was somewhat weaker during the early peak of the poststimulus BHA response in sensory ROIs. This is possibly due to a ceiling effect on BHA estimates driven by supra-threshold stimulation, or to the sparse electrode coverage of primary visual and auditory areas, preventing us from observing potential effects on early sensory responses.

By contrast, several previous studies have revealed a mixed pattern of results, showing a positive (Barry et al., 2000; Baslar & Stampfer, 1985; Becker et al., 2008; Dockree et al., 2007; Jasiukaitis & Hakerem, 1988; Mo et al., 2011; Roberts et al., 2014) or non-linear relationship (Kloosterman et al., 2019) between ongoing alpha power and other measures of poststimulus excitability (e.g., neuronal firing, non-invasive high-frequency power, and late ERP components). One possible explanation for these mixed results is that this relationship may depend on mechanisms other than functional inhibition. For example, ERPs are thought to reflect (1) stimulus-related neural activation, which is presumably suppressed during functional inhibition, and (2) baseline shift of non-zero-mean oscillations (Vadim V. Nikulin et al., 2007; Mazaheri & Jensen, 2008), which results in ERP amplification during states of strong prestimulus alpha power (Iemi et al., 2019). Accordingly, to rule out the possibility that the observed relationship between alpha oscillations and BHA was due to baseline shift, we estimated how BHA and ERPs were related to prestimulus alpha power separately for periodic sites with positive and negative oscillatory mean within the alpha band. We found that states of strong prestimulus *negative*-mean alpha oscillations were related to a more *negative* prestimulus ERP baseline and, in turn, a more *positive* poststimulus ERP signal. By contrast, states of strong prestimulus *positive*-mean alpha oscillations were related to a more *positive* prestimulus ERP baseline and, in turn, a more *negative* poststimulus ERP signal. In other words, states of strong prestimulus power were associated with a non-zero prestimulus ERP baseline, resulting in an amplification of the poststimulus ERP (in absolute value), consistent with baseline shift and with previous studies using non-invasive electrophysiology (Becker et al., 2008; Iemi et al., 2019). These results demonstrate that baseline shift contributes to the generation of the ERP (Vadim V. Nikulin et al., 2007). By contrast, BHA decreased with prestimulus alpha power regardless of the polarity of the oscillatory mean. Therefore, we conclude that the alpha-BHA relationship reflects an interaction between oscillatory activity and excitability consistent with functional inhibition, rather than a consequence of baseline shift.

Finally, it should be noted that, to maximize the number of trials for within-subject analysis, we used all trials regardless of whether or not the stimulus onset or content could be predicted based on the preceding cues (see Methods). While ongoing fluctuations in prestimulus activity before the (task-irrelevant) noise image reflect spontaneous changes in internal processes including attention, arousal, or motivation, fluctuations before the other (task-relevant) visual cues and the target syllable may also reflect changes in predictability. Future studies are necessary to differentiate how these different endogenous processes modulate ongoing neural activity and behavior.

### Relationship between prestimulus alpha power and behavior

Based on the functional inhibition account, we hypothesized that prestimulus alpha oscillations modulate task performance (e.g., RT in the auditory discrimination task). We found slower RTs following states of strong prestimulus alpha power estimated in the window before the task-relevant target stimulus (i.e., in syllable-locked epochs) across both periodic and aperiodic sites, including auditory and somatomotor regions. It is important to note that previous studies reported both positive (Bollimunta et al., 2008; Bompas et al., 2015; Kelly & O’Connell, 2013; Kirschfeld, 2008; Lou et al., 2014, 2014; Mazaheri et al., 2014; Min & Herrmann, 2007; Paoletti et al., 2019; van den Berg et al., 2016; Zhang et al., 2008), negative (Bollimunta et al., 2008; Del Percio et al., 2007; Zhang et al., 2008), and null relationships (Andino et al., 2005; van Dijk et al., 2008; Bays et al., 2015) between prestimulus alpha power and RT. This mixed evidence may be due to whether power is estimated in a brain region processing task-relevant or -irrelevant information, as well as to task differences (e.g., whether accuracy or speed is emphasized), different laminar organization across regions (Bollimunta et al., 2008, 2011; Mo et al., 2011), or to long-range temporal dependencies in both alpha power and RT estimates, resulting in spurious positive and negative correlations (Schaworonkow et al., 2015).

Interestingly, we observed that RT was positively related to alpha power in periodic sites, but also to the aperiodic signal in the same frequency band (i.e., alpha power in aperiodic sites). This may be explained by residual periodic activity in individual trials in aperiodic sites, or by a recent proposal (Donoghue et al., 2020; Peterson et al., 2018) suggesting that, in addition to alpha oscillations, the aperiodic signal modulates information processing and thus behavior (e.g., RT: Zhang et al., 2008).

### Interrelation between prestimulus alpha power, behavior, and excitability

Based on the functional inhibition account, we hypothesized that prestimulus alpha oscillations influence behavior by modulating excitability during stimulus processing. Therefore, key to our analysis was the mediation between prestimulus alpha power and behavior via poststimulus BHA (Judd & Kenny, 1981). We found that in periodic sites (1) single-trial prestimulus alpha power was negatively correlated with poststimulus BHA, and (2) positively correlated with RT (consistent with our binning analysis), (3) even after controlling for poststimulus BHA; and (4) that single-trial poststimulus BHA was negatively correlated with RT, (5) even after controlling for prestimulus alpha power. Critically, controlling for poststimulus BHA reduced the correlation between prestimulus alpha power and RT (i.e., indirect effect) in periodic sites (but not in aperiodic sites), suggesting that alpha oscillations and BHA explained a similar portion of RT variability. This is consistent with mediation and indicates that the behavioral changes associated with alpha oscillations (rather than the aperiodic signal in the same band) are related to the influence of alpha on excitability.

Notably, some previous studies attempted to examine the interrelation between alpha oscillations, excitability, and behavior (e.g., Hartmann et al., 2015; Kayser et al., 2016; Vugt et al., 2018). Some analyzed concurrent neural and behavioral changes as a function of ongoing/prestimulus alpha power, without directly assessing their interrelation (Bollimunta et al., 2008; Haegens et al., 2011; Min & Herrmann, 2007; Mo et al., 2011; Vugt et al., 2018). Other studies used statistical methods to directly test for said interrelation, but the results were null or inconclusive, possibly because excitability was estimated with indirect/non-invasive measures (stimulus-evoked response: Wöstmann et al., 2019; decoding metric: Kayser et al., 2016; McNair et al., 2019). Accordingly, to the best of our knowledge, our study is the first to show that a modulation of poststimulus excitability mediates the relationship between prestimulus alpha oscillations and behavior, consistent with functional inhibition. However, it is important to note that our results do not allow us to distinguish whether the mediation reflects a direct effect of alpha oscillations, or a consequence of an additional process affecting all variables in parallel. Future research should address this question by testing whether alpha oscillations directly cause the mediation effect, ideally by using neuromodulation techniques (Helfrich et al., 2014; Romei et al., 2010).

### Relationship between prestimulus alpha power and neural stimulus feature encoding

Based on the functional inhibition account, we hypothesized that prestimulus alpha oscillations affect how stimulus information is encoded in excitability measures (i.e., poststimulus BHA). Using BHA temporal patterns, we estimated decoder accuracy and confidence across different states of prestimulus alpha power and between periodic and aperiodic sites. We found that decoder confidence decreased with alpha power most prominently in periodic sites with a negative alpha-BHA relationship. This is consistent with the functional inhibition account and suggests that the BHA reduction during states of strong alpha power may underlie the decrease in decoder confidence. By contrast, across all sites, decoder accuracy was unaffected by prestimulus alpha power, even after controlling for overall decoder accuracy and for the alpha-BHA relationship.

How could the decoder indicate higher confidence without becoming more accurate during states of increased excitability? We speculate that weak prestimulus alpha power is associated with higher BHA responses, but also with proportionally more variability, leaving the signal-to-noise ratio (i.e., mean-to-variance ratio, or Fano factor) unchanged (Goris et al., 2014; Tolhurst et al., 1981; Tomko & Crapper, 1974). However, as mean and variance of the neural response increase, the neural stimulus representations fall farther from the classifier’s discriminant boundary or hyperplane (see model in Figure 5G). This modulation is expected to affect the overall strength of neural stimulus representations (i.e., decoder confidence) for both correct, but also incorrect trials, without affecting their precision (i.e., decoder accuracy). Future work is necessary to determine whether the effects of alpha oscillations on the magnitude of neuronal excitability are multiplicative (as assumed in our model) or additive, and whether they are proportional to changes in response variability, ideally by using single-unit recordings which enable an estimation of spike mean and variability.

Our findings that prestimulus alpha oscillations affect decoder confidence, not accuracy, are consistent with a growing number of studies reporting that strong prestimulus alpha oscillations are related to reduced subjective, rather than objective measures of perceptual decision-making in both detection (less liberal criterion: Iemi et al., 2017; Limbach & Corballis, 2016) and discrimination tasks (lower visibility: Benwell et al., 2017; lower confidence: Samaha, Iemi, et al., 2017). Future research is necessary to establish whether lower decoder confidence, reflecting decreased stimulus encoding strength, underlies the perceptual effects in these studies.

Previous studies have analyzed how decoding performance is related to alpha oscillations, though the results were inconsistent, showing positive (Kayser et al., 2016; McNair et al., 2019), negative (Barne et al., 2020; van Ede et al., 2018), and null relationships (Griffiths et al., 2019). In the studies reporting a positive relationship, stimulus features were decoded using low-frequency EEG activity (1–70 Hz), which includes alpha oscillations. Therefore, it is possible that high signal-to-noise ratio may result in strong alpha power estimates as well as higher decoding performance, potentially leading to a spurious positive correlation. To avoid such circularity, in other previous studies stimulus identity was decoded using the hemodynamic fMRI BOLD signal (Griffiths et al., 2019) or the low-frequency activity after removing alpha-band activity (van Ede et al., 2018). In our study, we addressed this issue by applying decoding analysis on poststimulus BHA, which does not include the alpha band, and reflects a more direct excitability measure than the fMRI signal. We reasoned that any relationship between alpha oscillations and decoding performance would reflect a genuine modulation of stimulus encoding, rather than a consequence of a potentially circular analysis.

It is important to note that we found no significant effect of prestimulus alpha power on decoder accuracy, consistent with one previous report using representational similarity analysis (Griffiths et al., 2019). By contrast, one previous study (van Ede et al., 2018) found that lower decoder accuracy (i.e., Mahalanobis distance) was related to strong prestimulus alpha power in posterior EEG, specifically in the presence of poststimulus distractors. When distractors were absent (as in our paradigm), decoder accuracy was no longer related to prestimulus alpha power, suggesting that the effect on decoder accuracy might emerge when the task requires the suppression of task-irrelevant information, which is believed to be supported by alpha oscillations (Haegens et al., 2010). Moreover, in another study (Barne et al., 2020), lower decoder accuracy (i.e., AUC) was related to strong prestimulus alpha power in parieto-occipital EEG electrodes when attention was cued to the spatial location of the to-be-decoded sensory stimuli. Therefore, it is possible that decoder accuracy may be affected by attention-induced/local (as opposed to spontaneous/global) fluctuations of alpha oscillations. Accordingly, future studies are necessary to determine whether decoder accuracy is influenced by a top-down modulation of alpha oscillations, ideally by using paradigms manipulating distractors and spatial attention.

### Conclusions

In sum, we demonstrate that strong prestimulus oscillations in the alpha+ band (i.e., alpha and neighboring frequencies), rather than the aperiodic signal, are associated with (1) decreased BHA (i.e., low neuronal excitability) before and after sensory input. Furthermore, we show that strong prestimulus alpha oscillations result in (2) slower perceptual decisions, and (3) reduced sensory encoding strength. These results provide a link between neural oscillations, excitability, and task performance, consistent with functional inhibition: we propose that, by modulating neuronal excitability, ongoing alpha+ oscillations affect behavior and neural stimulus representations.

## Methods

### Participants

This study involved nine individuals with medication-resistant epilepsy (5 females; mean age, 29 years, SEM=4). All patients had intracranial electrodes implanted as part of presurgical diagnosis of epilepsy. Data collection was performed at the Comprehensive Epilepsy Center of New York University Langone Health, and was approved by the Institutional Review Board at New York University Langone Health. Verbal and written informed consent were collected from all patients before participation in the study in accordance with the Declaration of Helsinki. Results from six individuals have been previously reported (Auksztulewicz et al., 2018).

### Experimental design

Each trial started with the presentation of a fixation cross (1.5 to 2 s), followed by a sequence of visual and auditory stimuli: a noise image, a picture of a scene, a picture of a face, and an auditory syllable (Figure 1A). The order of the sequence was fixed across trials. Each of the visual stimuli and the auditory stimulus were presented for a duration of 0.210 and 0.250 s, respectively. Visual stimuli were displayed at the center of a laptop screen placed at the bedside at approximately a 70-cm distance. The noise image consisted of grayscale random horizontal and vertical lines and was identical in all trials. For the other stimuli, different exemplars were presented across trials, selected from four different scene images (e.g., the White House or Taj Mahal), eight different face images (e.g., Barack Obama, George W Bush), and two different syllables (/ga/ vs. /pa/) The target syllables were produced by a male speaker; during the experiment, they were played with speakers at 70 dB or levels comfortable for the patient. The participants were asked to perform a 2-alternative-forced-choice discrimination of the target syllable (“which syllable did you hear: “/ga/ vs. /pa/?”) with a speeded button press using the index and middle fingers of the right hand. The experiment was implemented in Presentation (Neurobehavioral Systems; https://www.neurobs.com/).

The behavioral paradigm was based on a 2 × 2 factorial design with factors “when” predictability and “what” predictability, resulting in four different conditions, each of which was recorded in a separate run, and randomized across participants (for more information see Auksztulewicz et al., 2018). In brief, “when” predictability was manipulated by varying the timing between visual and auditory stimuli across trials: the four stimuli comprising a trial were presented with either a fixed inter-stimulus interval (ISI) of 1 s in the temporally predictable runs or with a randomly jittered ISI (mean 1 ± 0 to 0.5 s random jitter) across trials in the temporally unpredictable runs. In addition, “what” predictability was manipulated by using contingencies between the visual stimuli and the target syllable. In the content-predictable runs, specific sequences of scene/face images predicted the identity of the target syllable (75% probability), while in the content-unpredictable runs, the two target syllables were equiprobable across trials. To increase statistical power of single-trial analyses, we combined trials from all predictability conditions with a sufficient prestimulus window (i.e., trials with ISI >= 1s).

### Neural recordings

The electrodes consisted of 8 × 8 grids of subdural platinum-iridium electrodes embedded in Silastic sheets (2.3-mm-diameter contacts, Ad-Tech Medical Instruments) with a minimum 10-mm center-to-center distance implanted over the temporal/frontal cortices, with additional linear strips of electrodes and/or depth electrodes. In this dataset, grid, strip and depth-electrodes were used at 41, 43, 16% of the sites, respectively (total N = 1044 electrodes). Recordings were obtained using the Nicolet ONE clinical amplifier (Natus). During recording, the signal was bandpass filtered from 0.5 to 250 Hz, digitized at 512 Hz, and online referenced to a screw bolted to the skull.

We performed electrode localization using previously described procedures (Yang et al., 2012). In brief, for each patient, we obtained preoperative and postoperative T1-weighted MRIs, which were subsequently co-registered and normalized to an MNI-152 template, allowing the extraction of the electrode location in MNI space. We assigned anatomical labels to each electrode using FreeSurfer cortical parcellation (Fischl et al., 2004) based on the Desikan-Killiany atlas (Desikan et al., 2006), resulting in 983 localized electrodes out of 1044. The Freesurfer suite provides an automated labeling of the cerebral cortex into units based on gyral and sulcal structure.

Electrophysiological data analysis was performed using custom-built MATLAB code (version R2019a; The MathWorks; RRID:SCR_001622) and the Fieldtrip toolbox (version 2018.08.01; www.ru.nl/neuroimaging/fieldtrip). Continuous signals were notch-filtered at 58–62 Hz and harmonics using zero-phase Butterworth filters. The data were re-referenced to a common average and segmented into epochs from −1.5 to 1 s relative to the onset of each visual and auditory stimulus (stimulus-locked) and relative to the motor response (response-locked epochs). In total, we obtained 96(trials)*5(epochs)*4(runs) = 1920 epochs per patient.

We rejected artifactual electrodes in which the BHA time-course during the 1-s prestimulus window exceed 5 standard deviations from the mean in at least 50 trials. We corroborated this automatic procedure with visual inspection and removed a total of 76 artifactual electrodes across 9 patients.

We discarded 48 epochs per patient with an ISI <1s. Additionally, we rejected artifactual epochs in which either the raw signal or the BHA time-course (at any electrode) from −1 to 1 s relative to stimulus onset exceeded 25 standard deviations from the mean. We corroborated this automatic procedure with visual inspection and removed 136 artifactual epochs per patient. We analyzed 1595 epochs per patient (i.e., 344 noise-locked, 278 scene-locked, 281 face-locked; 356 syllable-locked; 334 response-locked epochs). For RT and single-trial analysis (mediation and decoding), we further discarded noise- and syllable-locked epochs with premature (RT<0.1 s; 10 epochs per patient), and late (i.e. RT > 0.8 s; 37 epochs per patient) RTs.

### Spectral Analysis

We computed power spectra for time windows before and after stimulus onset, separately for each electrode and for noise-, scene-, face, and syllable-locked epochs. The duration of the prestimulus window was 1 s for noise- and syllable-locked epochs, and 0.5 s (zero-padded to 1 for scene- and face-locked epochs to diminish the contamination due to event-related activity of previous visual stimuli. The duration of the poststimulus window was 1 s in all epochs. We multiplied these epochs with a Hanning taper, and estimated the spectra between 1 and 150 Hz (1-Hz frequency resolution) using a fast Fourier transform (FFT) approach.

We computed a single-trial estimate of the power in the alpha (7–14 Hz), beta (15–30 Hz), and BHA range (70–150 Hz). To avoid disproportionally representing power at the lower-bound frequencies (∼70 Hz) due to the 1/f property of the neural signal, we computed a frequency-normalized BHA by z-scoring the power spectra across trials per frequency. This normalization ensures that frequencies between 70 and 150 Hz equally contribute to the BHA estimates.

In addition, to inspect the time course of the spectral dynamics, we computed time-frequency representations (TFRs) of power. We estimated the low-frequency TFR (2–40 Hz; 1-Hz step size) using an adaptive sliding time window of three cycles length (Δt = 3/f), and the high-frequency TFR (70–150 Hz; 5-Hz step size) using a fixed window of 0.2 s, and applied a Hanning taper before estimating power using an FFT approach. The BHA time-course was computed by averaging the high-frequency TFR across frequencies.

### Spectral peak detection

To determine the individual alpha peak frequency, we detected the highest local maximum within the alpha range of the power spectra (using 1-s prestimulus noise-locked epochs) separately for each electrode (following the methods in Haegens et al., 2014). We refer to electrodes with and without alpha-band peaks as “periodic” and “aperiodic” sites, respectively. Note that the alpha range was set between 7 and 14 Hz in line with previous literature and to account for slower alpha oscillations previously reported in epilepsy patients (Stoller, 1949; Abela et al., 2019).

We used two complimentary methods to estimate beta-band peaks, both aimed to remove the 1/f (aperiodic) component of the power spectrum, as this obscures the smaller peaks in the beta band by strongly biasing lower frequencies. First, we used linear regression (least-squares fit) to fit a linear model to the log-transformed power spectra in the beta range. We subtracted the linear trend from the log-transformed power spectra and detected the beta-band peak as the highest local maximum within the beta range of the flattened power spectrum (Haegens et al., 2014; Nikulin & Brismar, 2006). Second, we detected beta-band peaks by estimating frequency regions within the beta range with power over and above the aperiodic 1/f signal using the fooof algorithm (Voytek et al., 2015; Donoghue et al., 2020) (modeled frequencies: 3–40 Hz, maximum number of peaks: 8, peak widths: 1–8, minimum peak height: 0.4 arbitrary units of power, peak threshold: 2 standard deviations; fixed approach). We refer to sites with and without beta-band peaks in both methods as “periodic” and “aperiodic” sites, respectively. Additionally, we identified a subset of beta-periodic sites without alpha-band peaks, which we refer to as “beta-only periodic” sites.

### Power-based binning analysis

To analyze the across-trial relationship between low-frequency oscillations and BHA, for each site we sorted trials into five bins (e.g., Linkenkaer-Hansen et al., 2004; Iemi et al., 2019) based on either alpha or beta power, separately for the pre- and poststimulus windows. For each bin we computed the average BHA for each time window. Similarly, we binned trials based on BHA, and computed the average TFRs of low- and high-frequency power spectra per bin. For group-level statistical analysis and visualization, the power spectra were normalized by the average power across all frequencies and bins, while the TFRs and BHA time-courses were normalized by the average power across time and bins per frequency.

### Statistical analysis

We used non-parametric cluster-based permutation tests (Maris & Oostenveld, 2007) for contrasts involving a temporal dimension. By clustering neighboring samples (i.e., time-frequency points) that show the same effect, this test controls for the multiple comparison problem while taking into account the dependency of the data. For each sample, a dependent-sample t-value was computed across sites for the relevant contrast (e.g., power difference between bins). We selected all samples for which this t-value exceeded an a priori threshold (p < 0.05), clustered these samples on the basis of temporal-spectral adjacency, and computed the sum of t-values within each cluster. By randomly permuting the data across the most extreme bins (bin 1 and 5) 1,000 times and determining the maximum t-sum on each iteration, we obtained a reference distribution of t-sums. A final p-value was calculated as the proportion of t-sums under the null hypothesis larger than the sum of t-values within clusters in the observed data. We adjusted the final alpha thresholds using Bonferroni correction for multiple comparisons (i.e., for multiple contrasts).

### Functional-anatomical regions of interest

We defined three experimentally relevant ROIs using a combination of functional and anatomical localizers. The functional localizer identified recording sites that were active during the presentation of the experimental stimuli (noise, scene, face, and syllable in stimulus-locked data) and during the behavioral response (in response-locked data). For each site we averaged the BHA time-course over the prestimulus window across stimulus-locked (−1.1 to −0.1 s) and response-locked epochs (−1.35 to −0.35 s). Then, for each site, we calculated the threshold as 2 standard deviations above the prestimulus BHA signal averaged across time points and trials. We identified stimulus-related and response-related sites as those sites whose poststimulus BHA time-course exceeded the site-specific threshold in stimulus-locked (0 to 0.5 s) and response-locked epochs (−0.25 to 0.25 s). Using this functional localizer, we identified 95 noise-related, 95 scene-related, 137 face-related, 300 syllable-related, and 565 response-related sites across 9 patients.

We complemented the functional localizer with an anatomical localizer. Specifically, we identified regions based on anatomical labels classically related to sensory processing and motor planning/response. Our visual localizer included the cuneus, lateral-occipital areas, lingual area, pericalcarine, fusiform gyrus, and inferior-temporal areas (N=123 sites). Our auditory localizer included transverse-temporal area, middle-temporal, superior-temporal, banks of the superior-temporal sulcus, supramarginal areas, pars opercularis, pars triangularis, superior-temporal gyrus (N=309). Our somatomotor localizer included precentral and postcentral areas (N=191).

Finally, we combined the functional and anatomical localizers to obtain the visual, auditory and somatomotor ROIs. Specifically, the overlap between visual anatomical sites and the combination of noise-, scene-, face-related functional sites yielded the visual ROIs (N=68). The overlap between the auditory anatomical sites and the syllable-related functional sites yielded the syllable-selective ROI (N=103), while the overlap between the somatomotor anatomical sites and the response-related functional sites yielded the somatomotor ROI (N=129).

### Baseline shift analysis

To determine whether baseline shift of non-zero-mean alpha oscillations contributes to the observed alpha-BHA relationship in periodic sites, we estimated the non-zero-mean property of alpha oscillations using the Baseline Shift Index (BSI: Vadim V. Nikulin et al., 2007, 2010) and the Amplitude Fluctuation Asymmetry Index (AFAI: Mazaheri & Jensen, 2008, 2010; van Dijk et al., 2010). In each patient and for each site, BSI and AFAI were estimated on the continuous (i.e., non-epoched) data and averaged across experimental runs. We classified periodic sites in two groups depending on whether BSI (or AFAI) within the alpha range was positive or negative.

To quantify BSI, the raw EEG data was first band-pass filtered using a 4^th^-order Butterworth filter centered at each frequency of interest ±1 Hz. Then, the Hilbert transform was used to extract a time-resolved power envelope. In addition, the raw EEG data was low-pass filtered using a 4^th^-order Butterworth filter with a 3-Hz cut-off frequency. We computed BSI as the Spearman correlation (ρ) between the power envelope and the low-pass EEG signal (i.e., slowly varying DC-like component) separately for each frequency and site (Iemi et al., 2019). When the low-pass signal is unaffected by power fluctuations, resulting in BSI = 0, there is evidence for zero-mean oscillations. Instead, when states of strong power result in positive (BSI>0) or negative (BSI<0) shifts of the low-pass signal, there is evidence for positive and negative oscillatory mean, respectively.

AFAI is based on the assumption that power fluctuations of non-zero-mean oscillations affect the peaks and troughs of the EEG signal differently. To quantify AFAI, we identified the time points of peaks and troughs as those data samples in the band-passed data which were larger (peaks) and smaller (troughs) than the two neighboring samples, respectively, and then estimated the magnitude of the raw EEG signal at these time points. We computed AFAI as the normalized difference between the variance at the time points of the peaks and troughs in the raw EEG signal separately for each frequency and site. Amplitude symmetry (i.e. AFAI = 0, power fluctuations equally modulate peaks and troughs) indicates a zero oscillatory mean. Amplitude asymmetry indicates a non-zero-mean: specifically, AFAI>0 indicates a stronger modulation of the peaks relative to the troughs, or a positive oscillatory mean; AFAI<0 indicates a stronger modulation of the troughs relative to the peaks or a negative oscillatory mean (see Nikulin et al., 2010 for a comparison between BSI and AFAI).

We used a binning approach to analyze how BHA in the prestimulus and poststimulus window changes as a function of prestimulus alpha power separately for the negative- and positive-mean sites. We repeated this analysis for ERPs: single-trial ERPs were computed on low-pass filtered data (<30 Hz) and baseline-corrected with the 1-s prestimulus signal.

### Behavioral analysis

We estimated behavioral performance using reaction times (RTs). Note that discrimination accuracy was at ceiling (∼90%) with only 36 incorrect responses per patient, thus preventing an analysis of trial-by-trial fluctuations. We used a binning approach to analyze how RTs on correct trials change as a function of prestimulus alpha power in periodic and aperiodic sites. We log-transformed RTs to correct for the skewness of their distribution. We averaged RTs across epochs for each bin, and normalized these estimates by the average RT across all epochs, separately for each patient. This analysis was run for noise- and syllable-locked epochs.

To understand the interrelation between prestimulus oscillations, behavior, and poststimulus excitability in syllable-locked epochs, we tested a mediation model in which prestimulus alpha oscillations (independent variable X) may affect RT on correct trials (dependent variable Y) via a modulation of poststimulus BHA (mediator, M). To this end, we used a causal step approach (Baron & Kenny, 1986; MacKinnon et al., 2000, 2007; Judd & Kenny, 1981) characterized by analyzing the correlation coefficients of four generalized linear models (GLMs).

The first GLM consists of a simple regression with independent variable predicting the mediator: *M = i + aX +e*, where *X* is prestimulus alpha power, *M* is poststimulus BHA, and *a* is the zero-order correlation coefficient reflecting the direct effect between *X* and *M*. In all GLMs, *i* refers to the intercept and *e* to the residual error of the model. Mediation requires that prestimulus alpha power is negatively correlated with poststimulus BHA (*c* < 0).

The second GLM consists of a simple regression with the mediator variable predicting the dependent variable: *Y = i + bM +e*, where *Y* is RT, *M* is poststimulus BHA, and *b* is the zero-order correlation coefficient reflecting the direct effect between *Y* and *M*. Mediation requires that poststimulus BHA is positively correlated with RT (*b* < 0).

The third GLM consists of a simple regression with the independent variable predicting the dependent variable: *Y = i + cX + e*, where *X* is prestimulus alpha power, *Y* is RT, and *c* is the zero-order correlation coefficient reflecting the direct effect between *X* and *Y*. Mediation requires that prestimulus alpha power is positively correlated with RT (*c* > 0).

The fourth GLM consists of a multiple regression with the independent variable and the mediator predicting the dependent variable: *Y = i + c’X + b’M + e*, where *X* is prestimulus alpha power, *Y* is RT, *M* is poststimulus BHA, and c’ and *b’* are the partial correlation coefficients reflecting the indirect effect between *X* and *Y* adjusted for *M*, and between *M* and *Y* adjusted for X, respectively. Mediation requires that poststimulus BHA is positively correlated with RT after controlling for prestimulus alpha power (*b’* < 0).

In addition, mediation requires a significant reduction in the effect of the independent variable on the dependent variable after accounting for the mediator, which can be estimated by the difference between zero-order and partial coefficients (c - c’): c > c’ (in absolute values) indicates partial mediation (or full mediation if c’ = 0). Note that mediation assumes that the independent variable X causes the mediator M (second GLM), and thus that the two variables are correlated. This correlation results in multicollinearity when the effects of independent variable and mediator on the dependent variable are estimated in a multiple regression model, yielding reduced coefficients in the fourth GLM.

We ran these four GLMs separately for each site, using single-trial FFT estimates of prestimulus alpha power, frequency-normalized poststimulus BHA, and RTs. Direct effects were estimated using variables that were normalized by z-scoring across trials, whereas the indirect effect was estimated using the original variables (i.e., not normalized). Note that, to compute unstandardized coefficients of the first and fourth GLM, we used RTs expressed in ms.

### Decoding analysis

We used multivariate pattern analysis (or “decoding”) to examine the relationship between prestimulus alpha oscillations and neural stimulus representations encoded in BHA. First, we tested whether stimulus features (i.e., syllable identity) could be decoded from the recorded neural activity; next we tested whether prestimulus alpha power modulates decoder accuracy and/or confidence.

We evaluated stimulus encoding in the time-course of neural activity using spatial decoding (similar to Gwilliams & King, 2020): we used l2-regularized logistic regression, under a stratified k-fold (k=4) cross validation scheme. Input features were normalized by the mean and the standard deviation of the training set. All decoding analyses were performed using the Python package scikit-learn (version 0.22.1: Pedregosa et al., 2011). The optimal regularization parameter was selected for each fold separately, by finding which of 10 log-spaced regularization strengths from 1e-4 to 1e+4 led to best model performance on the test set (using the LogisticRegressionCV scikit-learn function with default parameters). We used 1,000 maximum iterations of the ‘lbfgs’ optimization algorithm. The input features to the classifier were 17 time-samples (0 to 0.8 s relative to syllable onset) of either BHA (frequency-normalized estimate) or the low-passed signal (< 30 Hz; e.g., Gwilliams & King, 2020; King et al., 2016; Salti et al., 2015). We down-sampled the low-passed signal to match the temporal resolution of BHA. The categorical class labels corresponded to syllable identity (/ga/ vs. /pa/). We analyzed 310 trials for each patient, with 233 trials used for each training set and 76 for each test set. The model was trained and tested within each recording site separately, providing a prediction for each trial at each location over space, based on the multivariate pattern of activity over time.

We derived two decoding performance metrics from the classifier predictions: (1) AUC (i.e., decoder accuracy) and (2) the maximum probabilistic prediction (i.e., decoder confidence). For AUC, we evaluated the similarity between the true label categories of the test set and the probabilistic class labels (normalized distance from the fit hyperplane: “predict_proba” in scikit-learn) of the same trials. We computed the AUC under the null hypothesis by randomly shuffling the label categories to the classifier and obtained a p-value as the proportion of the null AUC estimates (across sites) that exceeded the true AUC independently for each site. We considered the AUC as significantly greater than chance if its p-value < 0.05. This analysis resulted in a decoding anatomical map indicating where syllable identity can be linearly decoded from temporal patterns of either BHA or low-passed signal (Figure 5 supplement). The maximum probabilistic prediction indicates the strength of the neural stimulus representation, regardless of whether or not it matches the ground truth; in other words, it is a measure of the classifier’s “confidence” about the true class label of a given trial, regardless of accuracy. When the probability = 0.5, the classifier’s predictions for the two class labels are identical (i.e., low confidence); when probability > 0.5, the classifier prediction for one class label is stronger than for the other class label (i.e., higher confidence).

We asked whether prestimulus oscillatory state shapes neural stimulus representations as reflected by decoding performance metrics. To test this, we evaluated how decoder accuracy and confidence based on BHA was related to trial-by-trial fluctuations of prestimulus alpha power. We used the classification procedure explained above, replacing the k-fold cross validation with a leave-one-out cross-validation (LOOCV) procedure. In LOOCV, the classifier is fit on all trials but one, evaluating model performance on the remaining “left-out” trial as a single-item test set. This is advantageous because: (1) it allows a maximal amount of data to be used for training, thus reducing noise in the model fit; (2) it provides a single-trial decoding estimate which can be analyzed by a binning approach. For each site and test trial, we computed the probabilistic estimates of the logistic regression for each syllable, grouping the results into five bins relative to prestimulus alpha power. Then, for each bin we estimated the AUC using the classifiers’ probabilistic estimates of the test trials and the vector of true class labels. Additionally, we normalized the maximum probabilistic estimates by z-scoring across trials, and computed decoder confidence by averaging these estimates within each bin.

## Acknowledgments

This work was supported by a grant from the Netherlands Organization for Scientific Research (NWO 016.Vidi.185.137, to S.H.), by the Silvio O Conte Center for Active Sensing (P50 MH109429, to L.I., C.E.S., S.H.), by the Basic Research Program of the National Research University Higher School of Economics (to V.V.N.), by the European Commission’s Marie Skłodowska-Curie Global Fellowship (750459 to R.A.), by a grant called Finding A Cure for Epilepsy and Seizures (FACES, to O.D.), by a NIH grant (1R01EB019805 to T.T), by the NYU Abu Dhabi Institute (G1001 to L.G.), and by the William Orr Dingwall Dissertation Fellowship (L.G.), by a re-entry supplement (R01DA038154, to Y.M.C).

We thank Esra Al, Julio Rodriguez Larios, Elie Rassi, and Hesham Elshafei for helpful comments on the manuscript.

---

**Figure 1 supplement.**
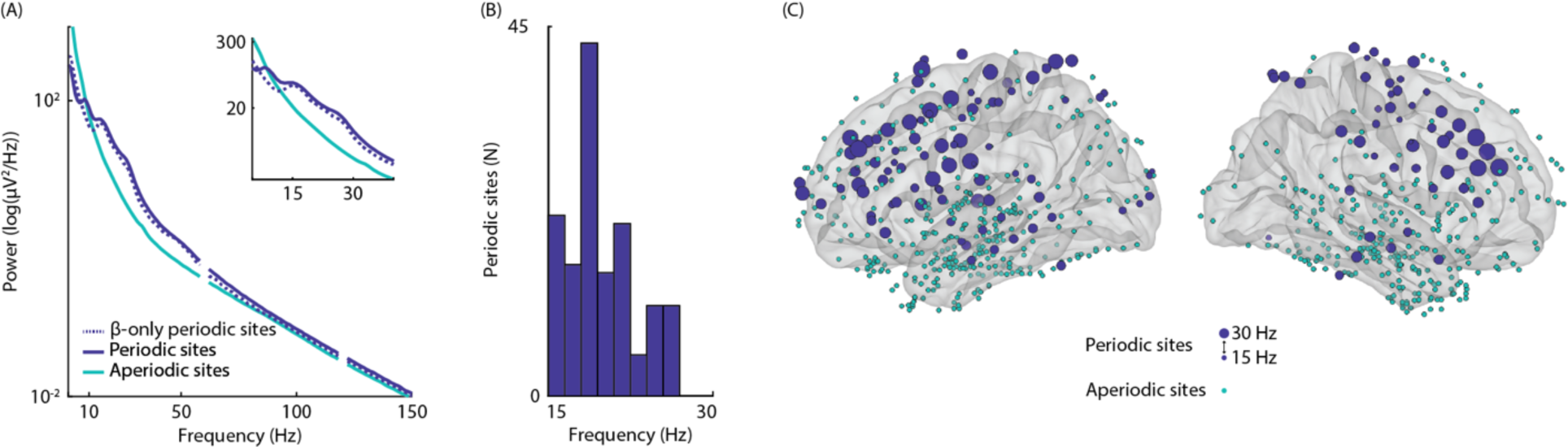
Beta-band periodic and aperiodic activity. **A.** Averaged power spectrum of the 1-s window before the noise image, shown separately for sites with beta oscillations (periodic sites, in blue), sites with beta oscillations but no detectable alpha-band peaks (beta-only periodic sites, blue dotted line), and sites without a detectable beta-band peak (aperiodic sites, in green). The power spectrum of periodic (and beta-only periodic) sites contains a beta-band peak (15–30 Hz) while the power spectrum of aperiodic sites shows only 1/f activity in the beta band. The inset shows the power spectrum for the frequency window of interest. **B.** Histogram of the beta-band peak frequencies of the prestimulus power spectrum across periodic sites. **C**. Schematic illustration of the iEEG electrode coverage. Blue and green dots illustrate periodic and aperiodic sites, respectively; the size of blue dots indicates the beta-band peak frequencies of periodic sites.

**Figure 2 supplement.**
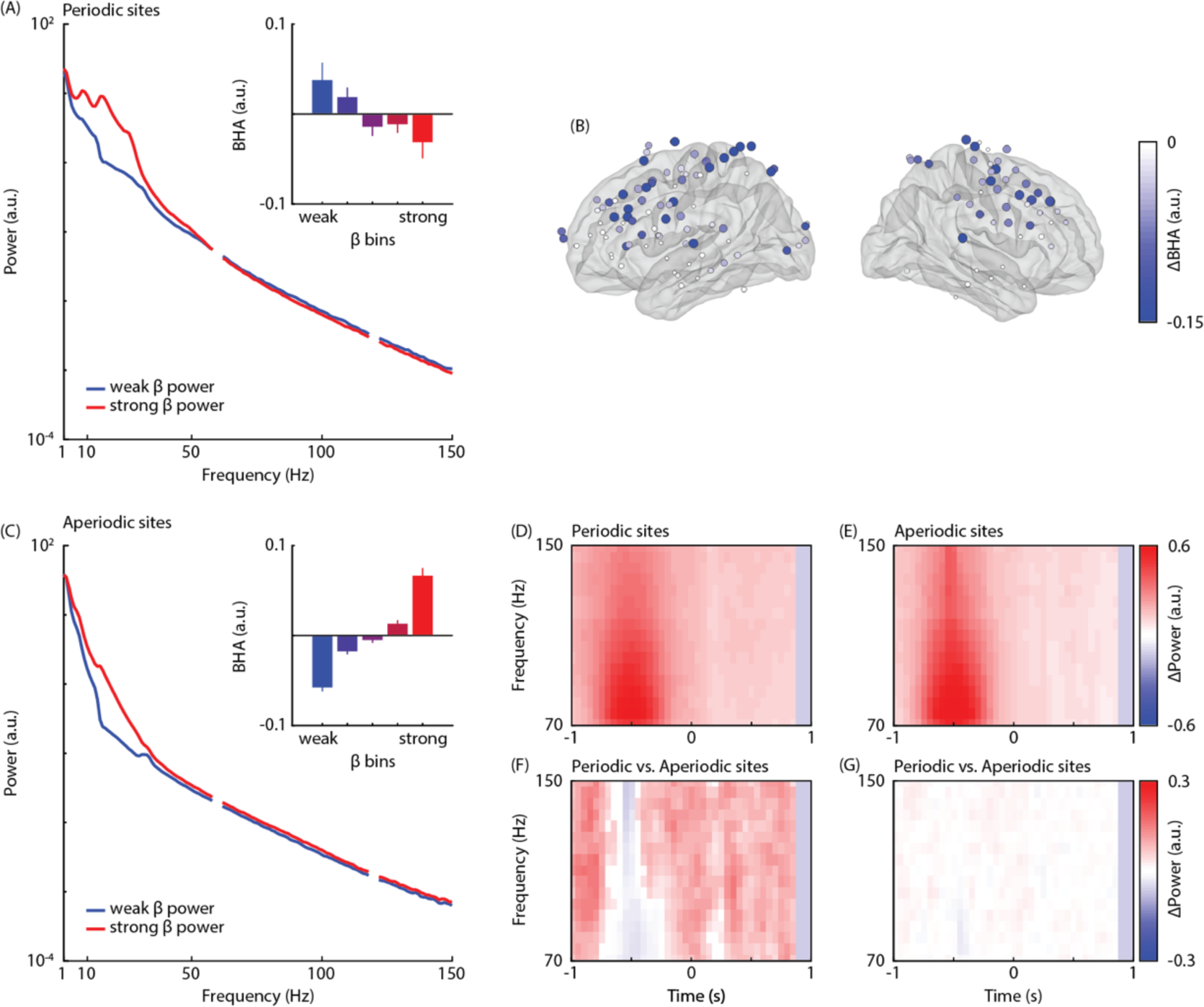
Correlation between prestimulus beta power and prestimulus BHA. **A.** Averaged power spectrum computed during the 1-s prestimulus window in noise-locked epochs, shown separately for bins of strongest (red) and weakest (blue) prestimulus beta power for periodic sites (spectra normalized with average power). The inset shows the averaged BHA (70–150 Hz) separately for five bins sorted from weakest (blue) to strongest (red) prestimulus beta power (normalized with the average across bins) for periodic sites. BHA decreases with beta power in periodic sites, consistent with functional inhibition. **B.** Anatomical map of the relationship between prestimulus beta power and BHA power in periodic sites, estimated as the normalized difference in BHA between bins of strongest and weakest beta power. The color and size of the dots are proportional to the experimental effect. The map shows negative effects (BHA in bin 5 < BHA in bin 1). No significant positive effects were found. **C.** Same as (A) for aperiodic sites, respectively. BHA increases with beta power in aperiodic sites, consistent with a broadband signal offset. In (A–C) periodic and aperiodic sites were defined relative to the beta band. **D/E.** Time-frequency representations of the difference in high-frequency power between bins of strongest and weakest BHA estimated during the prestimulus window in noise-locked epochs (normalized by average power across the most extreme bins and masked to show significant effects only) in periodic and aperiodic sites, respectively. The clusters indicate the spectral and temporal extent of the binning based on prestimulus BHA (note that this is a circular analysis for visualization purposes). Red clusters indicate the spectral and temporal extent of BHA-based binning. **F/G**. Time-frequency representations of the comparison of the high-frequency power difference between prestimulus BHA bins across periodic and aperiodic sites (normalized by average power across sites and masked to show significant effects only). Panels F and G show the results for all sites and a subset of periodic and aperiodic sites with matched high-frequency power difference between states of strong and weak BHA, respectively. Positive clusters (red) indicate that high-frequency power difference between BHA states is more positive in periodic sites. No negative clusters were found. In (D–G) periodic and aperiodic sites were defined relative to the alpha band.

**Figure 3 supplement.**
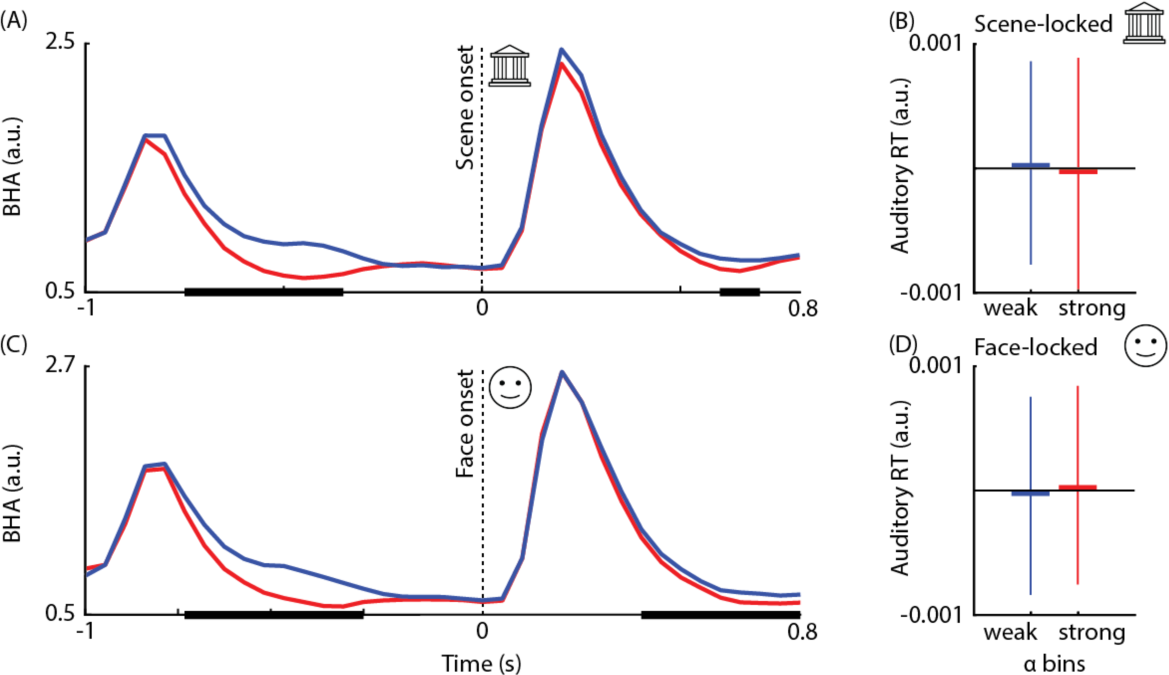
Correlation between prestimulus alpha power, poststimulus, and reaction times in the visual ROI during scene- and face-locked epochs. **A.** BHA time-course shown separately for bins of weakest (blue) and strongest (red) prestimulus alpha power for scene-locked epochs in the visual ROI. Bold horizontal black lines indicate significant differences based on cluster permutation testing. We found two significant negative clusters, indicating that states of weak prestimulus alpha power (i.e., before scene image onset) coincided with weaker prestimulus BHA (t=−33.32, p<0.001; from −0.75 to 0.35 s) and were followed by weaker poststimulus BHA (i.e., after scene image onset; t= −9.28, p=0.038, from 0.6 to 0.7 s), consistent with functional inhibition. **B.** Averaged RT, shown separately for the weakest (blue) and strongest (red) bin of prestimulus alpha power in scene-locked epochs in the visual ROI. RT is not affected by alpha power in visual areas estimated before the scene image (p>0.05). **C.** Same as in (A) for face-locked epochs. We found two significant negative clusters, indicating that states of weak prestimulus alpha power (i.e., before face image onset) coincided with weaker prestimulus BHA (t=−41.84, p<0.001; from −0.75 to 0.30 s) and were followed by weaker poststimulus BHA (i.e., after face image onset; t=−33.46, p<0.001, from 0.4 to 0.85 s), consistent with functional inhibition. **D.** Same as in (B) for face-locked epochs. RT is not affected by alpha power in visual areas estimated before the face image (p>0.05).

**Figure 5 supplement.**
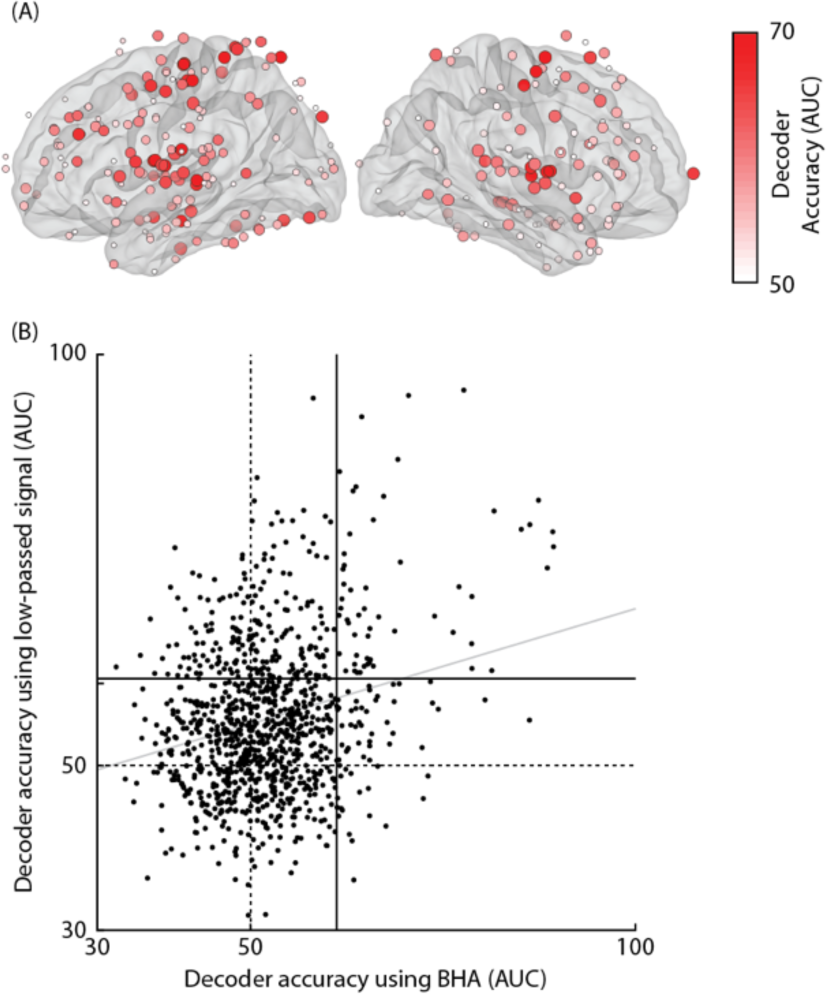
Relationship between prestimulus alpha power and neural stimulus decoding using BHA or low-passed signal. **A.** Anatomical map of decoder accuracy (AUC) indicating where syllable identity can be linearly decoded from BHA temporal patterns. We found 117 sites with an AUC significantly higher than chance (p<0.05): 23 sites belonged to our functional-anatomical auditory ROI, including the superior temporal gyrus, supramarginal gyrus, banks of the superior-temporal sulcus, and 15 sites belonged to our somatomotor ROI, including postcentral and precentral areas. Overall, the probability that the AUC was greater than chance (p-value AUC) did not differ between periodic and aperiodic sites (unpaired t(1042)=1.12, p=0.264). This demonstrates that syllable identity can be decoded using BHA temporal patterns, consistent with previous studies using other metrics of decoder accuracy (mutual information: Belitski et al., 2008; Peters et al., 2017), and using other decoding input (low-passed signal: King et al., 2016; Gwilliams & King, 2020). Note that, because the response mapping used in this study was fixed across trials, it is possible that decoding performance was based on the syllable-specific motor plan, rather than on stimulus information per se. However, we found significant AUC in both the somato-motor and auditory ROIs, suggesting that the decoder was successful in using not only motor but also sensory information. In the figure, red dots illustrate sites where the classifier successfully predicted syllable identity (AUC>0.5). The size and color intensity proportionally reflect the AUC magnitude. (A) includes only periodic sites with AUC>0.5. **B.** Across-site correlation between the AUCs obtained using low-passed signal and BHA temporal patterns. In addition to BHA, we estimated AUC using the low-passed signal (<30 Hz) as an input to the decoder and found a significant positive correlation with the estimates obtained using BHA (Spearman rho=0.12 p<0.001), suggesting that stimulus information contained in BHA was comparable to that obtained using a brain signal that has been commonly used in previous studies (Gwilliams & King, 2020; King et al., 2016; Salti et al., 2015). In the figure, each dot represents the AUC values obtained from spatial decoding based on BHA (on the x axis) and on the low-passed signal (on the y axis). The dotted lines at 0.5 represents theoretical chance AUC, while the bold lines represent the empirical chance AUC based on permutation test. The grey line indicates the least-squares fit. (B) includes both periodic and aperiodic sites.

**Figure 6 Supplement.**
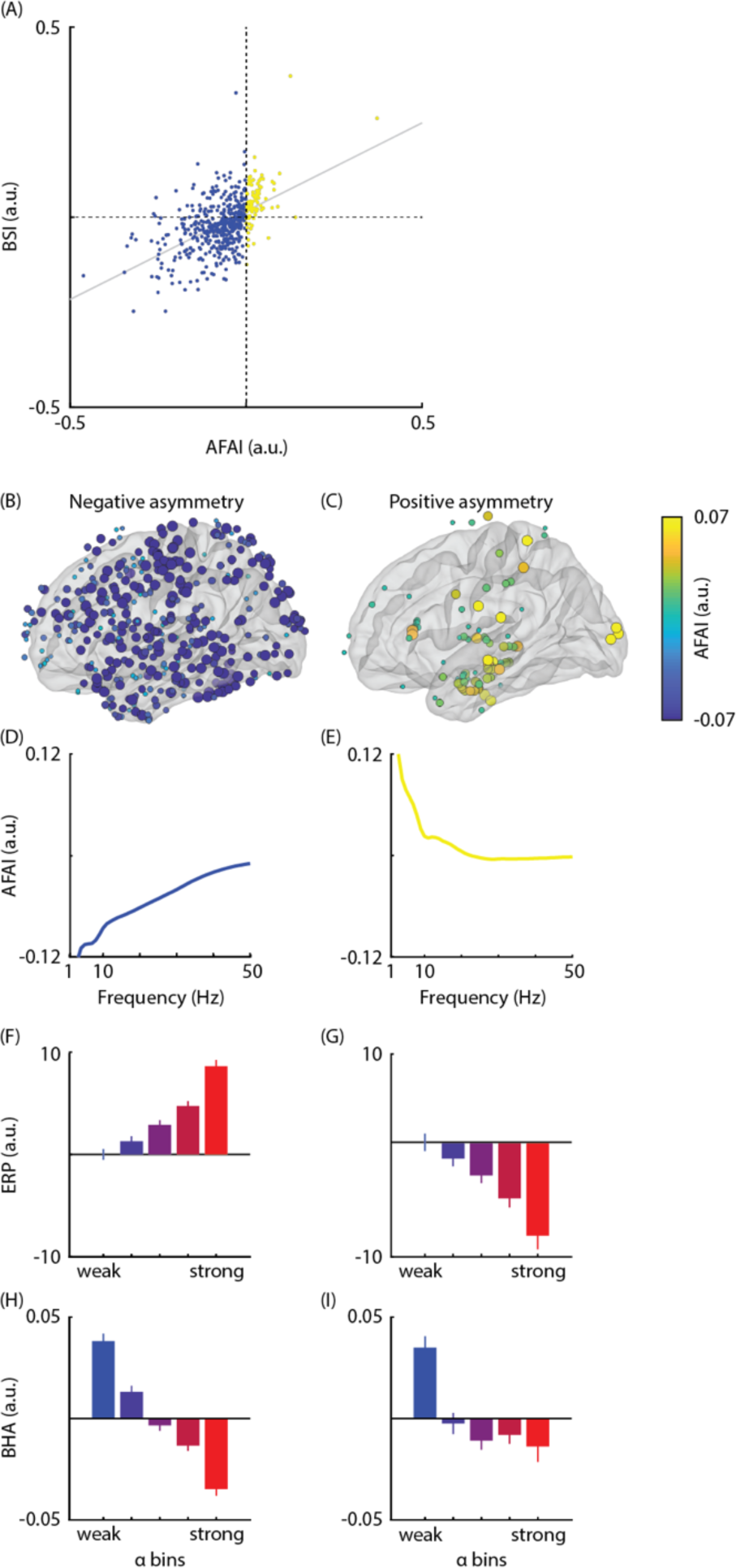
Correlation between alpha power and prestimulus BHA accounting for amplitude asymmetry. **A.** Relationship between the magnitude of the baseline-shift-index (BSI; y axis) and amplitude fluctuation asymmetry index (AFAI; x axis). There is a moderate positive correlation between BSI and AFAI across sites (Spearman rho = 0.54 p<0.001). **B–I.** Same as in Figure 6 (C–J) but the split between sites is based on the polarity of the amplitude asymmetry, indexed by AFAI, which is correlated with the polarity of the oscillatory mean. The AFAI-based analysis replicates the BSI-based analysis reported in the main text.

## Notes

### Competing Interest Statement

The authors have declared no competing interest.

